# Altered RNA-processing provides a mechanistic framework delineating human sex-reversal associated with pathogenic variants in the RNA-helicase DHX37

**DOI:** 10.1101/2025.01.10.632330

**Authors:** Maëva Elzaiat, Estelle Talouarn, Somboon Wankanit, Laurène Schlick, Caroline Eozenou, Joëlle Bignon-Topalovic, Etienne Kornobis, Pierre-Henri Commere, Chloé Baum, Valérie Seffer, Ken McElreavey, Anu Bashamboo

**Affiliations:** Human Developmental Genetics, CNRS UMR3738, Institut Pasteur, Université Paris Cité, Paris, France; Department of Pediatrics, Faculty of Medicine Ramathibodi Hospital, Mahidol University, Bangkok, 10400, Thailand; Institut Pasteur, Université Paris Cité, Plate-forme Technologique Biomics, F-75015 Paris, France; Institut Pasteur, Université Paris Cité, Bioinformatics and Biostatistics Hub, F-75015 Paris, France; Cytometry Platform, Institut Pasteur, Université Paris Cité, Paris, France; UTechs Single-Cell Biomarkers Platforms, Institut Pasteur, Université Paris Cité, Paris, France

**Keywords:** DHX37, 46, XY DSD, sex-determination, HyperTRIBE, single-cell full-length transcriptomics, induced Sertoli-like cells

## Abstract

Recurrent heterozygous missense variants in the highly conserved RNA-helicase *DHX37,* which is required for ribosome biogenesis, are a frequent cause of 46,XY sex-reversal or testis regression syndrome. How these missense variants specifically disrupt testis formation is unknown. Here, we demonstrate that mutant DHX37 proteins retain their ATPase activity and are not associated with stabilization of cellular β-catenin. Transfection of DHX37 p.R674Q mutant protein in an *in-vitro* cellular model recapitulating human Sertoli cell formation, showed a reduced activation of pro-testis genes compared to the WT protein. The expression of a DHX37 mutant protein in *in-vitro* derived human Sertoli-like cells (iSLCs) was also associated with global changes in gene expression, predicted to impact basic cellular functions. To define RNA transcripts interacting with either the WT or a mutant (p.R674Q) protein, we combined HyperTRIBE and single-cell full-length RNA-sequencing approaches using iSLCs. Gene ontology analysis indicated that transcripts targeted by WT DHX37 were primarily associated with cytoskeleton organization, including cell motility and cell adhesion. However, in contrast transcripts targeted by the mutated DHX37 protein, were not only associated with cytoskeleton organization but also with protein degradation and cell death. These data provide mechanistic framework that may explain how variants in the DHX37 protein can result in 46,XY sex-reversal through altered RNA networks that are required for the formation and maintenance of the supporting cell lineages of the human testis.

## Introduction

Sex-determination, where bipotential somatic precursors develop as either Sertoli (testis) or granulosa (ovaries) cells, is the key process in gonad formation. Commitment to either of those fates is the outcome of a battle between poorly conserved and characterized, mutually antagonistic male and female gene regulatory networks (GRNs) that canalize development down one organogenetic pathway, whilst actively repressing the alternate^1^. In mammals, the Y-linked Testis-Determining Factor SRY (Sex-determining Region of the Y chromosome) upregulates its downstream target *SOX9* (SRY-box transcription factor 9) beyond a critical threshold^2^. Following this, SOX9 autoregulates and initiates a positive feedforward loop with FGF9 Fibroblast Growth Factor 9)/FGFR2 (Fibroblast Growth Factor Receptor 2)^3–5^ and PGDS (Prostaglandin D2 Synthase)/PGD2 (Prostaglandin D2)^6–8^ to maintain and amplify its expression. During differentiation, fetal Sertoli cells proliferate, aggregate and surround the spermatogonia to form the seminiferous cords, characterized by epithelialization of Sertoli cells^9^. In XX individuals, in the absence of SRY, the -KTS isoform of WT1 is required to initiate ovary formation^10^. Further granulosa cell differentiation is regulated by two independent GRNs; RUNX1/FOXL2 (Runt-related transcription factor 1/Forkhead box L2)^11,12^ and WNT4/RSPO1 signaling to activate β-Catenin^13,14^. Stablization of β-Catenin in XY gonads disrupts testis-determination and results in male-to-female sex-reversal^15^.

Disruption of GRNs regulating human gonad development results in a continuum of pathologies termed Disorders/differences of sex development (DSD), where there is discordance between the chromosomal, anatomical and physiological sex. 46,XY individuals with gonadal DSDs are characterized by feminization or under-virilization due to complete (CGD) or partial gonadal dysgenesis (PGD)^16^. 46,XY testicular regression syndrome (TRS), includes absence of gonadal tissue on one or both sides in association with atypical genitalia (MIM 273250). Familial occurrences of 46,XY CDG/PGD and TRS^17,18^ suggest a common underlying genetic etiology for these group of pathologies.

Recurrent heterozygous missense variants in *DHX37* (DEAH-Box Helicase 37) gene are a frequent cause of 46,XY gonadal dysgenesis and TRS^19–22^ (reviewed in ^23^). DHX37 is a ubiquitously expressed RNA-dependent ATPase, whose structure and function are conserved through to yeast. The protein is composed of two tandemly repeated RecA-like domains, RecA1 and RecA2, that include eleven structural motifs involved in specific protein functions including, ATP binding and hydrolysis (I, II and VI), RNA binding (Ia, Ib, Ic, IV, IVa, V) and coordination of ATP hydrolysis and unwinding (III and Va)^24^. Biallelic variants in DHX37 are associated with a distinct autosomal recessive disorder termed NEDBAVC (Neurodevelopmental disorder with brain anomalies with or without vertebral or cardiac anomalies) without any genital anomalies (MIM 618731)^25^. The two groups of variants, associated either with DSD or NEDBAVC, are located within or immediately adjacent to the functional domains of the protein^23^. However, to date, no common variant has been identified for these two distinct pathologies. The individuals presenting with PGD/CGD and TRS do not have any associated somatic anomalies^19^. This raises the question of how the mutations in a ubiquitous factor, with a highly conserved structure and function, can cause a phenotype restricted to gonadal anomalies?

DHX37 is indispensable for ribosome biogenesis in eukaryotic cells^26,27^. Most of our knowledge of the biological function of DHX37 has come from the analysis of its yeast orthologue, Dhr1^28^. An early step in the maturation of the small subunit (SSU) of the ribosome is regulated by a megacomplex, called SSU processome, that is formed by the association of ribosomal proteins, RNA-binding factors and rRNA precursors^29^. A core element of the SSU processome is the small nucleolar RNA U3 (U3 snoRNA) required for facilitating interactions within the megacomplex^29,30^. During ribosome biogenesis, Dhr1/DHX37 interacts with and sequesters U3 snoRNA from the megacomplex^26,27^. Consistent with this, Dhr1 mutations in yeast or inhibition of *DHX37* in HeLa cells impairs the maturation of the 18S rRNA, leading to reduced levels of 40S subunits^26,27^. This suggests that the role of DHX37 in ribosome biogenesis is conserved from yeast to human cells.

Along with its key role in ribosome biogenesis, other cell/tissue specific role for DHX37 have been reported. Genome-wide CRISPR screens identified DHX37 as a regulator of anti-tumor properties in CD8 T cells^31^. In mouse and human T cells, DHX37 was shown to suppress CD8 T cell activity, including lymphocyte activation, cytokine production, regulation of cell-cell adhesion or interferon-gamma production by interacting with components of the NF-κB pathway^31^. In hepatocellular carcinomas, DHX37 was shown to promote liver cancer cell proliferation and cell-cycle progression^32^ by interacting with PLRG1 (Pleiotropic regulator 1) to transcriptionally activate cyclin D1 expression. In the zebrafish, Dhx37 physically interacts with GlyR α1, α3, and α4a subunit transcripts and regulates specific splicing events^33^. Fish carrying a homozygous missense mutation, p.K489P, exhibit abnormal escape behavior characterized by atypical dorsal bend before swimming. In the mutant fish, the expression levels of GlyR α1, α3, α4a, and βa subunits were decreased due to splicing defects, resulting in abnormal motor response caused by a deficit in glycinergic synaptic transmission^33^. However, the quantities of ribosomal RNA precursors remained unchanged between the wild-type and mutant fish. These data provide evidence for additional biological functions for DHX37, independent of ribosome biogenesis.

We attempted to address the mechanism(s) by which mutations in *DHX37* can affect differentiation/development of somatic cells in the 46,XY gonad. We show that DHX37 sex-reversing mutant proteins retain ATPase activity and they do not induce nucleolar stress or stablization of β-catenin. Introduction of the sex-reversing p.R674Q mutation in *in-vitro* derived human Sertoli-like cells (iSLCs)^34^ resulted in a reduced activation of pro-testis gene expression. To further determine how variants in DHX37 might disrupt human testis-development, we defined RNA transcripts targeted by the wild-type or pathogenic p.R674Q protein in iSLCs using a combination of HyperTRIBE and single-cell full-length RNA-seq^35^. Transcript analysis indicated that the DHX37 mutants may affect testicular development by dysregulating multiple biological processes including, but not limited to, cytoskeleton organization (including cell motility and cell adhesion), protein degradation and cell death. This preliminary data provides a framework that can help us to understand the mechanism(s) associated with mutations in *DHX37* and 46,XY DSD.

## Methods

### Cell culture

#### Induced Sertoli-like cells culture

Induced Sertoli-like cells (iSLCs) were derived from human 46,XY iPSC line as previously described^34^. Cells were cultured at 37°C with 5% CO_2_ and maintained in Supporting medium [Advanced DMEM (#12634010, ThermoFisher Scientific) supplemented with 5 mL of 100X Insulin, Transferrin, Selenium (#12097549, ThermoFisher Scientific) and 50 µL of Epidermal Growth Factor (20 ng/mL; human recombinant #ab9697, Abcam)] with a change every 2-3 days.

#### siRNA-mediated silencing

Knockdown (KD) experiments were performed using 65,000 sorted iSLCs^34^, which were seeded into 6-well plates the previous day to reach 50% confluence at the time of transfection. Cells were transiently transfected with 30 pmoles of siRNAs against *DHX37* (#4392420, Silencer^TM^ Select Pre-Designed siRNAs: ID# s33511, s33512 and s33513, ThermoFisher Scientific), *ACTN4* (#E-011988-00-0020, Accell Human ACTN4 siRNA-SMARTpool, Dharmacon, Horizon Discovery), *CDC42* (#E-005057-00-0020, Accell Human CDC42 siRNA-SMARTpool, Dharmacon, Horizon Discovery), *PTK2* (#E-003164-00-0020, Accell Human PTK2 siRNA-SMARTpool, Dharmacon, Horizon Discovery), or an equal amount of the Silencer^TM^ Select Negative Control No. 1 siRNA (#4390843, ThermoFisher Scientific) using Lipofectamine RNAiMax (ThermoFisher Scientific) following the manufacturer’s instructions. Six hours after transfection, culture medium was added, and cells were allowed to grow for 48 hours before performing subsequent experiments. The KD efficiency was verified at the transcript and protein levels. A total of three independent experiments was performed.

#### Plasmids and site-directed mutagenesis

The expression vector containing wild-type human *DHX37* full-length cDNA coding sequence (NM_032656) fused to DDK tag (pCMV6-Entry, #RC207812) was purchased from OriGene (Rockville, Maryland, États-Unis). For HyperTRIBE experiments, DHX37 was fused to ADAR2^E488Q^ (Adenosine deaminases acting on RNA) hyperactive-catalytic domain (AA 299-701) required for the conversion of adenosine into inosine by deamination^36^. A 50 AA rigid linker made of 10 repetitions of EAAAK peptide was designed to link *DHX37* and *ADAR2cd(E488Q)* sequences. Briefly, *DHX37* full-length cDNA sequence was extracted from OriGene expression vector using the forward primer 5’-CGCTAGCAGATCTGCCACCATGGGGAAGCTGCGC-3’ and the reverse primer 5’-CAGCCTCCTTTGCCGCTGCCTCCTTAGCTGCCGCTTCCTTCGCGGCAGCTTCTTTAGCTGCGGCCTCCTTTGCAGCTGCCTCTTTAGCTGCAGCTTCGTGGACAGTGGTGGGGGGCCAG-3’. *ADAR2cd(E488Q)* sequence was extracted from CIRTS-8: bdefensin 3-TBP6.7-hADAR(299-701)E488Q vector (#132545, Addgene) using the forward primer 5’-CGCGAAGGAAGCGGCAGCTAAGGAGGCAGCGGCAAAGGAGGCTGCTGCGAAAGAAGCCGCAGCCAAGGAGGCGGCGGCTAAAGAAGCTGCATAAATTGCACTTGGATCAGACGCCA-3’ and the reverse primer 5’-CGTCGACG AATTCTCAGGGCGTGAGTGAGAACT-3’. The two PCR fragments were fused by overlap extension PCR and the final amplicon was introduced into the pIRES2-acGFP1 vector (#642435, Takara Bio) using the restriction enzymes *BglII* and *EcoRI*. The pathogenic c.2021C>A (p.Arg674Glu, p.R674Q) variant was introduced in the vectors described above using the QuikChange II Site-Directed Mutagenesis Kit (# 200524, Agilent) with the forward primer: 5’-CGAGCGGGCAGAGCAGGACAGACGGAGCCCGGCCACTGCTACAG-3’ and the reverse primer 5’-CTGTAGCAGTGGCCGGGCTCCGTCTGTCCTGCTCTGCCC GCTCG-3’ following the supplier’s recommendations. The pIRES2-acGFP1 vector containing only the *ADAR2cd(E488Q)* sequence was used as a negative control. It was obtained after isolation of the insert using the forward primer 5’-GATCCGCTAGCAGATCTATGATGTTGCACTTGGATCAGACGCCA-3’ (containing an ATG) and the reverse primer 5’-CGTCGACGAATTCTCAGGGCGTGAGTGAGAACT-3’ and introduction of it in the pIRES2-acGFP1 vector using *NheI* and *EcoRI* restriction enzymes. Plasmids were amplified following heat-shock transformation of NEB-5alpha competent cells (#C2992H, BioLabs), and purified with the NucleoBond Xtra Maxi Plus kit (#740416, Macherey-Nagel). The sequence of all plasmids was confirmed by direct sequencing before performing functional studies.

#### Over-expression of WT and p.R674Q DHX37

The effect of WT and p.R674Q mutant proteins on gene expression was assessed after over-expression of WT or p.R674Q DHX37 in iSLCs. Briefly, 65,000 iSLCs were seeded into 6-well plates and transfected with 500 ng of wild-type or pathogenic variant-containing pCMV6-DHX37-DKK expression vector, using FuGENE 6 (#E231A, Promega) as the carrier (µL):DNA (µg) ratio of 3:1. Cells were lysed 48 hours after transfection.

For the HyperTRIBE experiments, iSLCs were transfected as described above with pIRES2-DHX37-ADAR2^E488Q^cd-GFP (pIDA) containing the WT or mutant DHX37, or the negative control pIRES2-ADAR2^E488Q^cd-GFP (pIA). Cells were lysed 48 hours after transfection and prepared for FAC sorting.

### FAC sorting

The pIRES2-DHX37-ADAR2^E488Q^cd-GFP construct produces the fusion protein and GFP from the bicistronic transcript. Therefore, iSLCs transfected with HyperTRIBE constructs were sorted using GFP fluorescence. Fluorescent cells were collected in FACS tubes with cell strainer (#352235, Corning) and examined using MoFLO Astrios with Summit v62 (Beckman Coulter, France) at 25 PSI with a 100 nM nozzle at approximately 2000 events per sec. GFP fluorescence was read with the 488 laser (576/21 band pass). After positive selection for Alexa 488, cells were collected in Eppendorf tubes containing 200 µL of supporting medium. On an average, GFP positive cells represented 8-13% of the initial population (**Supp Fig. S1**) and cell death was about 6-8%. Between 30,000 (pIDA: WT, p.R674Q) and 40,000 (pIA) sorted cells were centrifuged and 10,000 cells were used for single cell cDNA library preparation.

### Single-cell cDNA library preparation for full-length RNA-sequencing

#### cDNA preparation for single-cell (10X genomics)

The GFP-positive sorted single-cell suspension was treated for “Chromium Next GEM Single Cell 3’ v3.1 protocol” (CG000315 Rev C, 10x genomics) in accordance with the manufacturer’s instructions, for cDNA preparation. Single-cell cDNAs were purified using 0.8X AMPure XP beads (Beckman Coulter, USA). Quantification was then performed with a Qubit fluorometer (Thermo Fisher Scientific, USA) and quality was determined using a Bionalyzer (Agilent Technologies, USA).

#### ONT library preparation and sequencing

Libraries were prepared following the “Single-cell sequencing on PromethION” protocol (V10x-SST_v9148_v111_revC_12Jan2022-promethion), with a modification to enable sample multiplexing. Specifically, the PCR-cDNA Barcoding kit (SQK-PCB111.24, Oxford Nanopore Technologies) was used instead of the cDNA-PCR Sequencing kit (SQK-PCS111). The resulting barcoded libraries were validated by both Qubit fluorometer and Bioanalyzer as described above. Sequencing was performed on PromethION flowcells (FLO-PRO002) using a P2 solo sequencer (Oxford Nanopore Technologies) for a duration of 72 hours. Each sample was loaded onto a dedicated flowcell. Data were base called and demultiplexed with MinKNOW version 22.07.7 using Guppy version 6.3.9.

### Identification of RNA editing events in single-cell RNA-seq data

Fastq files were first processed using BLAZE^37^ and FLAMES^38^ in order to obtain count matrices. These count matrices were then used to obtain the barcodes of cells which expressed at least one copy of *DHX37* or *ADARB1*. The fastq files were then filtered to keep only the reads found in these cells. We aligned the reads on the hg38 assembly using Minimap2^39^. Bam files were sorted and duplicates were marked. Variants were called using FreeBayes^40^ and annotated using SnpEff^41^. Variants were then filtered according to the following criteria: (1) A>G variants in transcripts encoded by the forward strand and T>C in transcripts encoded by the reverse strand, (2) QUAL > 20, (3) depth > 10, (4) read ratio of at least 20% for the alternative allele, (5) no significant Fisher strand bias, (6) variant not previously identified in the WGS of the cells. For every gene, we kept the variants annotated for the canonical/MANE transcript. Variants found in the pIA (negative control) were removed from “WT” and “p.R674Q” samples. For subsequent analysis, we analyzed transcripts for which at least three variants were found.

### Differentially Expressed Genes analysis

The count matrix of each sample was obtained using FLAMES^38^ and processed independently to remove cells with low numbers of transcripts, low numbers of UMIs as well as high percentages of mitochondrial and ribosomal genes. Doublets were removed and the expression of each transcript were regressed on the cell cycle scores and the expression of *DHX37* and *ADARB1* were used as markers of transfection efficiency. All files were then integrated using Seurat^42^.

### Gene pathway enrichment analysis

Gene ontology analysis was performed on the differentially expressed genes and edited transcripts using STRING Database (v12.0)^43^ focusing on the biological process (BP) gene ontology terms. For the edited transcripts, they were separated in two different populations: the ones edited in presence of WT DHX37 (n=785) and the ones specifically edited in presence of p.R674Q DHX37 (n=397). Cytoscape (v3.10.2, https://cytoscape.org/)^44^ was used to display networks.

### RNA extraction and RT-qPCR

Total RNA was extracted from sorted iSLCs using TRIzol reagent (#15596026, ThermoFisher Scientific). RNA yield was quantified with a NanoDrop spectrophotometer (NanoDrop Technologies), and 1000 ng of RNA was used to synthesize complementary DNA (cDNA) with the Quantitect Reverse Transcription Kit (#205311, QIAGEN) following manufacturer’s recommendations. cDNA was diluted 1/25 prior to the qPCR. qPCR was performed using TaqMan Universal Master Mix II, with UNG (#4440038, Applied Biosystems) on a StepONEPlus qPCR machine (Applied Biosystems). The following TaqMan probes (Applied Biosystems) were used; *ACTN4*: #Hs00245168_m1; *AMH*: #Hs00174915_m1; *BRACHYURY/T*: #Hs00610080_m1; *CDC42*: #Hs00918044_g1; *CTNNB1*: #Hs00355049_m1; *DHX37*: #Hs01553956_m1; *DMRT1*: #Hs00232766_m1; *FGF9*: #Hs00181829_m1; *FOXL2*: #Hs00846401_s1; *GATA4*: #Hs0171403_m1;; *MDM2*: #Hs00540450_s1; *NANOG*: #Hs02387400_g1; *PTK2*: #Hs01056457_m1; *RPL19*: #Hs02338565_gH; *RSPO1*: #Hs00543475_m; *RUNX1*: #Hs01021970_m1; *SOX9*: #Hs01001343_g1; *TP53*: #Hs01034249_m1; *WNT4*: #Hs01573505_m1; *WT1*: #Hs01103751_m1. Relative mRNA levels were determined by calculating 2^-ΔΔCT^ values relative to the 18S rRNA normalizer gene (*RPL19*). Relative gene expression is presented using Microsoft Excel (Redmond, Washington, USA) or GraphPad Prism version 10.4.0 (GraphPad Software, Boston, Massachusetts USA, www.graphpad.com) as the mean 2^-ΔΔCT^ values, the reference sample (SCR or Empty vector (EV)) being set to 1. Pairwise comparisons were performed using the Wilcoxon test (Rstudio). A P-value of less than 0.05 was considered statistically significant.

### Protein extraction and Western Blot

Forty-eight hours after transfection/knock-down, cells were rinsed with 1X PBS and lysed with IP lysis buffer (#87788, ThermoFisher Scientific) supplemented with 100X Halt Protease Inhibitor Cocktail (#78440, ThermoFisher Scientific) and 100X EDTA (#78440, ThermoFisher Scientific) for 30 min. The lysates were centrifuged for 15 minutes at 20,000g and the supernatants were retrieved. Protein quantification was performed using the Pierce Detergent Compatible Bradford Assay kit (#23246, ThermoFisher Scientific) according to the manufacturer’s instructions. For Western Blot experiments, 10 µg of protein extracts were used for over-expression assays whereas 25 µg were used for the knock-downs. After a denaturation step using 4X XT loading buffer (#1610791, Bio-Rad) at 95°C for 5 minutes, protein samples were separated on Criterion XT 10% polyacrylamide gel (#3450112, Bio-Rad) and transferred to PVDF membrane (#T831.1, Merck Millipore). Membranes were blocked in Tris-buffered saline containing 0.1% TWEEN 20 (#27949, SIGMA) (TBS-T) and 5% non-fat powdered milk for an hour at room temperature. Incubation with primary antibodies was performed overnight at 4°C under stirring. Anti-DHX37 antibody produced in rabbit (1:4000, #HPA047607, Sigma-Aldrich), anti-beta Catenin Polyclonal Antibody (CAT-15) (1:2000, #71-2700, ThermoFisher Scientific) or WNT4 Recombinant Superclonal Antibody (9HCLC) (1:2000, #710889, ThermoFisher Scientific) were diluted in TBS-T containing 5% BSA. Membranes were then washed thrice (15 minutes each) with 1X TBS-T, and they were incubated with Goat anti-Rabbit IgG antibody coupled to the HorseRadish Peroxidase (HRP) (#ab205718, Abcam) diluted 1:15,000 in TBS-T containing 0.5% BSA for one hour at room temperature under stirring. The revelation was performed using Pierce ECL Western blotting substrate (#32132, ThermoFisher Scientific) and X-ray films. The detection of several proteins on the same blot was obtained by treating the membrane with Antibody Stripping solution (#L7710A, Interchim) followed by probing with new primary antibodies. β-Actin (1:2500, #A2228, Sigma-Aldrich) was used for normalization. Band intensity was quantified using Fiji software^45^, and results of quantification were plotted as the mean ± SEM of three independent experiments. The statistical comparison of the means was performed using a Student test.

### Malachite green phosphatase assays

To determine the ATPase activity of the different DHX37 mutant proteins, HEK-293T cells were transfected to express the different mutations and after 48h, protein extraction was performed as described above. The ATPase assay was performed using the malachite green method for the detection of released phosphate (#MAK307, SIGMA), following the supplier’s recommendations. After 0, 15, 30, 45 and 60 min of colour development, absorbance was measured at 620 nm, using the Glomax Multi+ Detection System (Promega). A standard phosphate curve was prepared (as per manufacturer’s instructions) in the enzyme buffer solution to determine the amount of free phosphate released by the enzyme variants tested. Experiments were repeated three times. Student’s t-test was used to determine statistical significance.

### Immunofluorescence

For immunostaining, iSLCs were cultured on Nunc™ Lab-Tek™ Chamber Slide systems (#177445PK, ThermoFisher Scientific). Forty-eight hours after transfection/knock-down, cells were rinsed with PBS and fixed using 4% PFA. Permeabilization was performed with 0.5% Triton X-100 (in PBS), followed by blocking of non-specific epitopes using PBS containing 5% BSA. Cells were then incubated with the primary antibody diluted in 3% BSA-PBS in a humid chamber overnight at 4°C. The following dilutions of primary antibodies were used: anti-DHX37 (1:200, #HPA047607, Sigma-Aldrich) and anti-beta Catenin Polyclonal Antibody (CAT-15) (1:500, #71-2700, ThermoFisher Scientific). Cells were then washed three times with PBS for 5 min. This was followed by incubation with the secondary antibody, either Goat anti-Rabbit IgG (H+L) Secondary Antibody, Alexa Fluor® 594 conjugate (#A-11037, Life Technologies) or Goat anti-Rabbit IgG (H+L) Secondary Antibody, Alexa Fluor® 488 conjugate (A-11008, Life Technologies) diluted 1:1000 in PBS with 3% BSA for 1h at room temperature away from light. After three washes, cells were incubated with DAPI for 15 min in the dark (1:2000, #62248, ThermoFisher Scientific). After three washes with PBS, slides were mounted using ProLong® Gold Antifade Mountant with DAPI (#P36931, ThermoFisher Scientific). Images were obtained with a Leica Microsystems DMI4000B microscope at 40x, 63x and 100x (with oil) magnifications or confocal ZEISS LSM 800 microscope and 20x or 40x oil objectives with an optical sectioning in Z every 1 μm and a tile scan of 10 to 15 Z stacks. Image analyses were performed with Fiji software^45^. The HyperTRIBE constructs in pIA or pIDA vectors were transfected as described previously into HEK-293T cells for validation. In addition to DHX37, GFP detection was performed using the anti-GFP antibody (1:500, #ab13970, Abcam) and Goat anti-Chicken IgY (H+L) Secondary Antibody, Alexa Fluor™ 488 (1:1000, #A-11039, ThermoFisher Scientific).

## Results

### Pathogenic DHX37 proteins retain ATPase activity

Since the sex-reversing variants are located within the REcA1 and RecA2 domains of the DHX37 protein that are required for ATP-binding, ATP-hydrolysis as well as RNA-binding (**Figure 1A**)^23^, we tested the hypothesis that these variants may affect the ATPase activity of the protein. Using the colorimetric method to detect ATP hydrolysis by measuring the release of inorganic phosphate (**Figure 1B**), we found that the kinetics of phosphate release was similar between the WT and mutant proteins. Therefore, it seems unlikely that the loss of DHX37 ATPase activity is responsible for the sex-reversing phenotypes.

**Figure 1:**
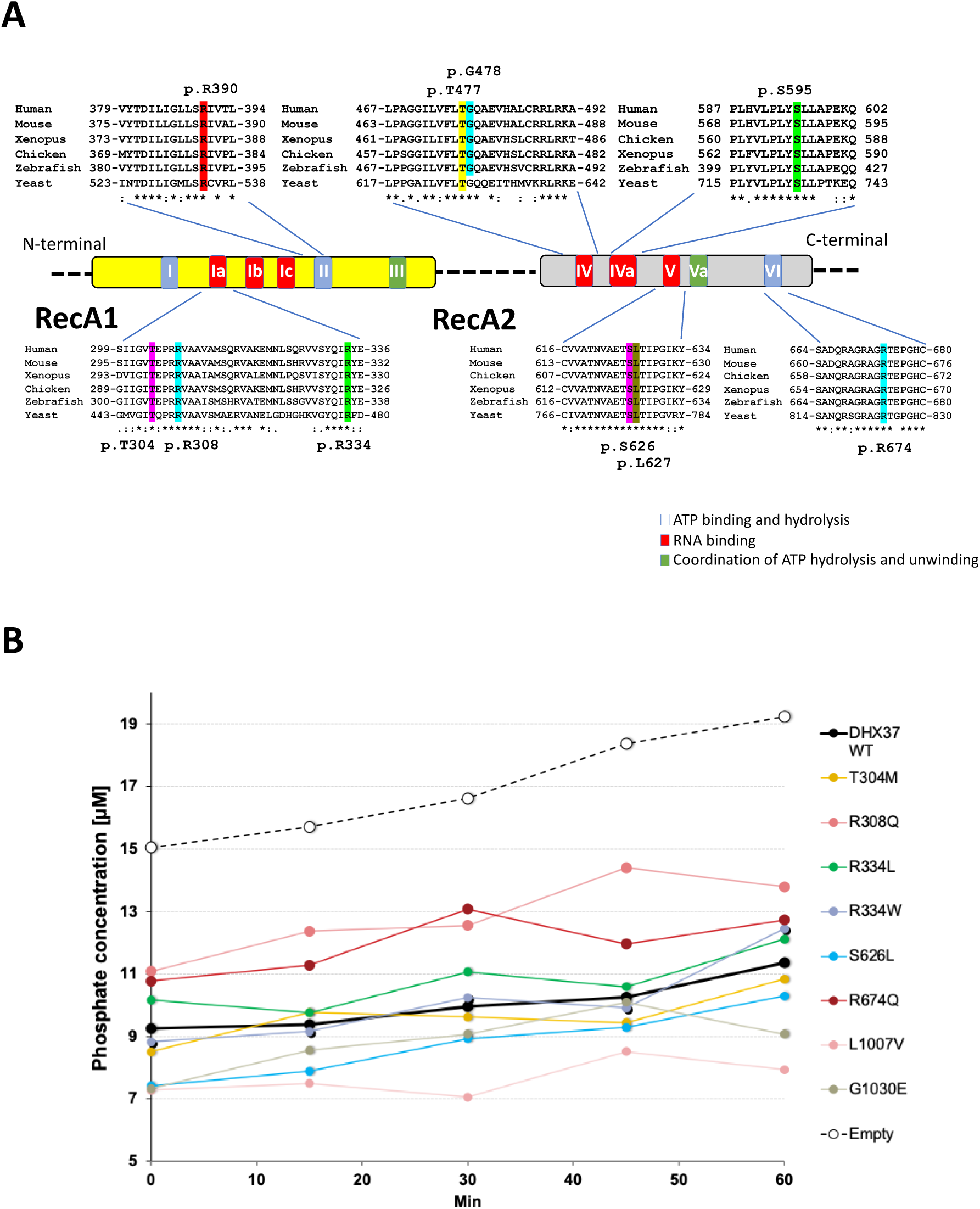
Pathogenic variants of DHX37 are not impaired for their ATPase activity. **(A)** Schematic structure of DHX37 protein. DHX37 is composed of two tandemly repeated RecA-like domains, containing eleven functional motifs. The amino acids involved in male-to-female sex-reversal or testicular regression syndrome are indicated. All are located within functional motifs and involve residues conserved from human to yeast. **(B)** Malachite Green colorimetric analysis of ATPase activity of DHX37 mutants. A green complex is formed when Malachite green molybdate reacts with inorganic phosphate under acidic conditions. Experiments were performed at constant protein and substrate concentrations while varying the incubation times. The plot shown is representative of three experiments. Statistical significance of the mean difference with respect to WT DHX37 for each time point was performed with the Student t-test.

### Testicular dysgenesis due to DHX37 variants does not result from β-Catenin stabilization

Pro-ovarian Wnt/β-Catenin signaling is a regulator of ribosome biogenesis^46^ and β-Catenin stabilization following nucleolar stress may serve as a compensatory mechanism to sustain ribosome biogenesis^47^. We hypothesized that the testicular dysgenesis associated with DHX37 mutations could be due to nucleolar stress induced stabilization of β-Catenin. Stabilization of β-Catenin is a known cause of male-to-female sex-reversal in the mouse^15^. To address this hypothesis, we knocked-down the endogenous *DHX37* or we overexpressed DHX37 WT or p.R674Q mutant proteins in iSLCs (**Figure 2**). Despite a KD efficiency of 80% at both the transcript and protein levels, we did not observe any significant change in β-Catenin expression, nor its distribution on the plasma membrane (**Figure 2A, B, C**). Similarly, we did not observe any significant change in β-Catenin expression, nor its distribution on the plasma membrane despite an average upregulation of 20,000 fold for WT or p.R674Q transcripts and a 5 fold increase in protein levels (**Figure 2D, E, F**). Moreover, neither the lack of DHX37 nor the overexpression of DHX37 WT and p.R674Q mutant, influenced the expression of the nucleolar stress markers such as *MDM2* and *TP53* (**Supp. Fig. S2**). These results suggest that DHX37 disruption in iSLCs does not lead to nucleolar stress and subsequent β-Catenin stabilization.

**Figure 2:**
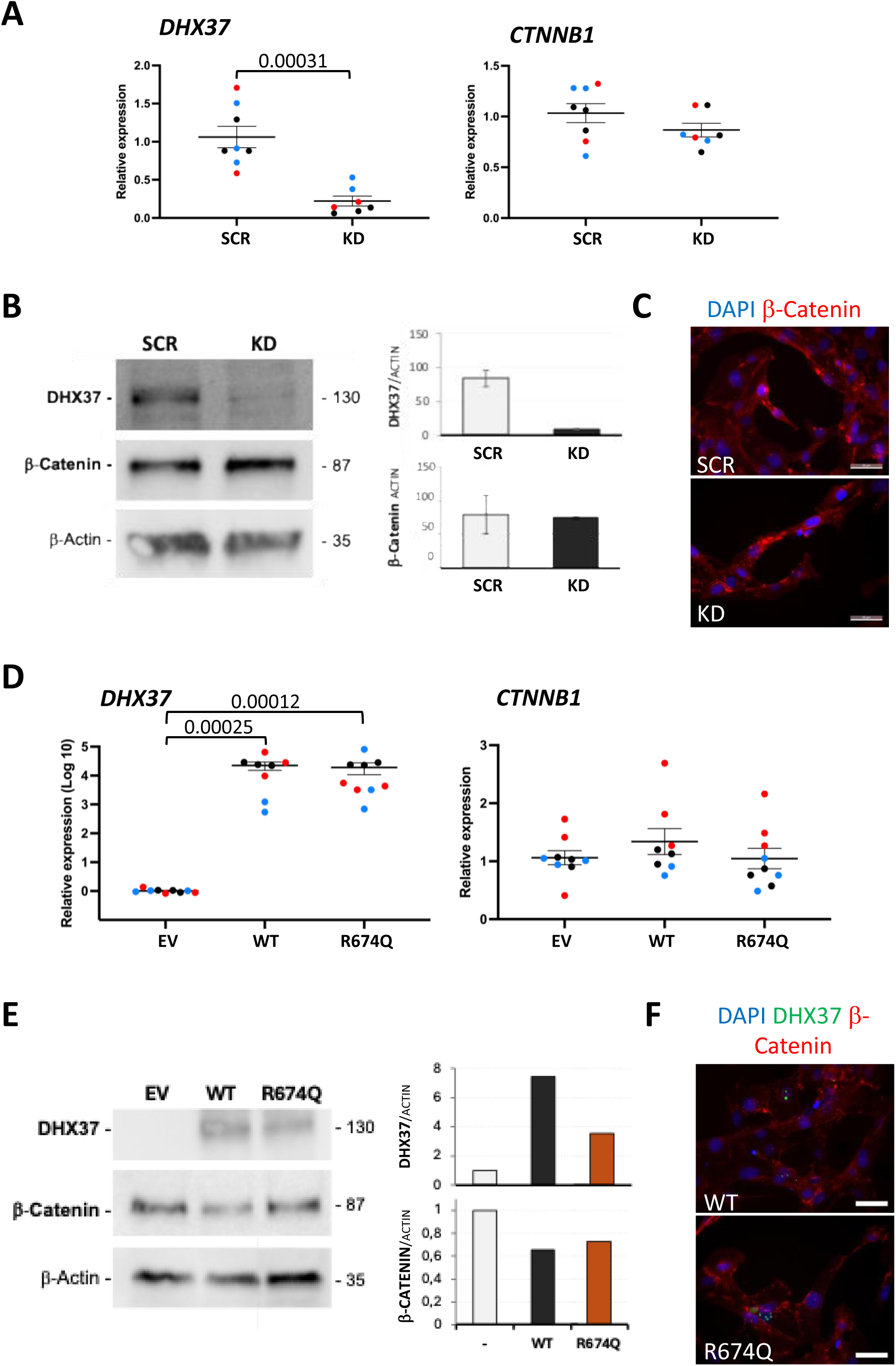
Disruption of DHX37 does not result in β-Catenin stabilization. **(A)** Effect of *DHX37* knock-down on *CTNNB*1 expression in iSLCs. The level of expression of endogenous *DHX37* and *CTNNB1* in the scramble (SCR) or knock-down (KD) condition was evaluated by RT-qPCR in three independent experiments. Expression was normalized against the *RPL19* gene and data plotted as ΔΔC_T_ values using GraphPad Prism. Pairwise comparisons were performed using the Wilcoxon test. A *p*-value of less than 0.05 was considered statistically significant. **(B)** Effects of *DHX37* knock-down on β-Catenin production in iSLCs. Production of endogenous DHX37 and β-Catenin in the SCR or KD condition was evaluated by Western Blot. β-Actin was used to normalize. Quantification of band intensities was performed with Fiji software. **(C)** Distribution of β-Catenin in iSLCs in the SCR or the KD was evaluated by immunofluorescence (IF). Scale bar: 50 µm. **(D)** The effect of overexpression of DHX37 WT and R674Q variants on *CTNNB*1 expression in iSLCs was evaluated by RT-qPCR as described above. **(E)** The consequence of overexpression of DHX37 WT and R674Q variants on β-Catenin production in iSLCs was evaluated by Western Blot. β-Actin was used for normalization. Quantification of band intensities was performed with Fiji software. **(F)** Distribution of β-Catenin after overexpression of WT or R674Q DHX37 in iSLCs in assessed by IF. Scale bar: 50 µm.

### In iSLCs, p.R674Q induces a reduced expression of pro-testis genes, as compared to that by WT DHX37

We determined whether transient (48h) over-expression of DHX37 WT or p.R674Q mutant in iSLCs following transfection had an impact on the expression of key pro-testis (*GATA4, WT1, SOX9, AMH)* or pro-ovary genes (*RUNX1, FOXL2, RSPO1* and *WNT4* (**Figure 3**)). Introduction of WT DHX37 results in increased expression of *GATA4* (fold change: 9.04, *p*=0.006), *WT1* (fold change: 2.08, *p*=0.02), *SOX9* (fold change: 3.03, *p*=0.009), *RUNX1* (fold change: 2.26, *p*=0.006), *FOXL2* (fold change: 2.68, *p*=0.009) and *WNT4* (fold change: 6.20, *p*=0.006). However, the introduction of the p.R674Q mutant resulted in a reduced upregulation of the Sertoli cell-markers *SOX9* (fold change: 1.63) and *AMH* (fold change: 1.5), an increased expression of *FGF9* (fold change: 2.4) as well as a decreased expression of the granulosa cell-marker *WNT4* (fold change: -1.81).

**Figure 3:**
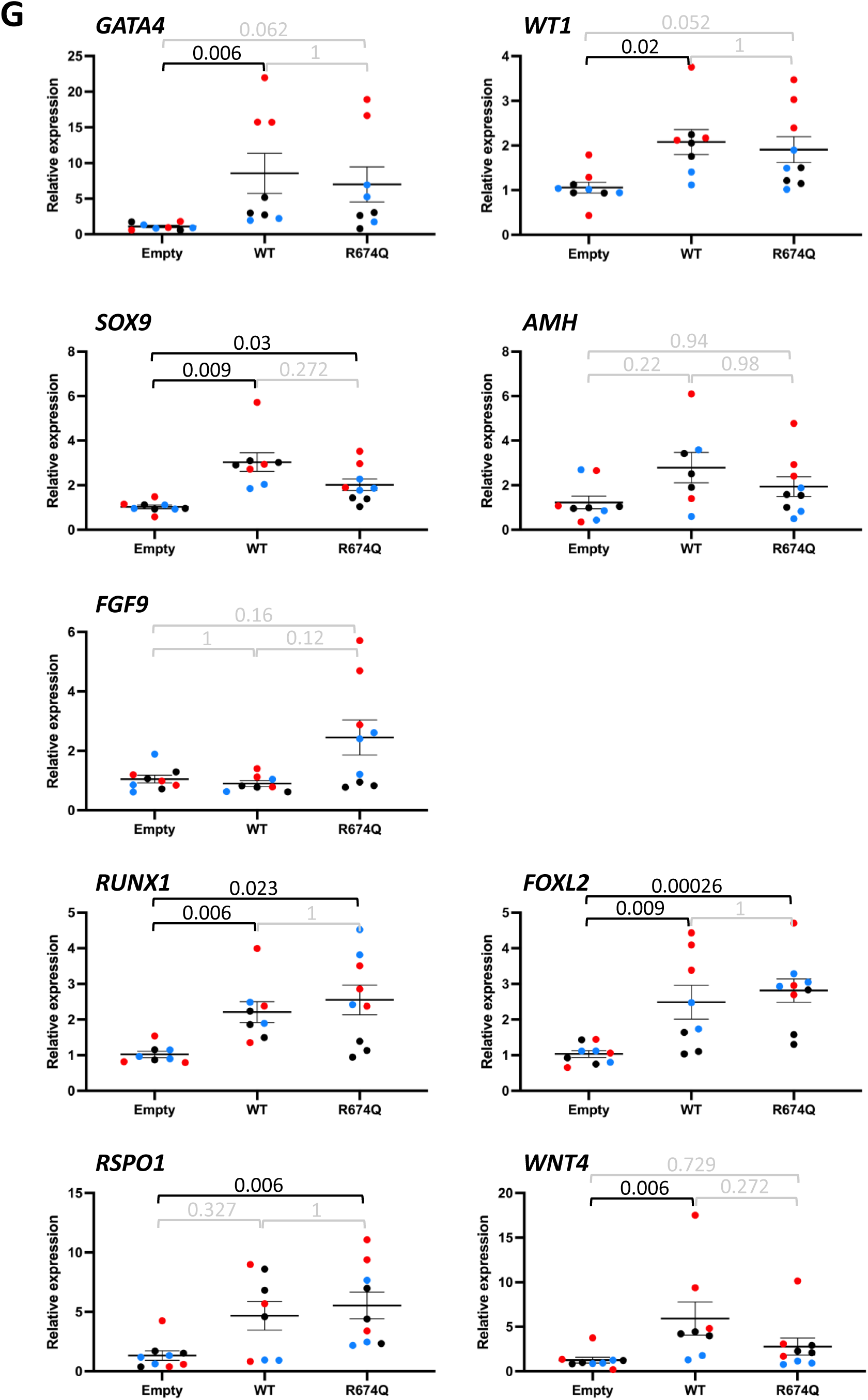
Overexpression of DHX37 WT and p.R674Q mutant protein influences sex-determining gene expression in iSLCs. The level of expression of *GATA4, WT1*, *SOX9*, *AMH*, *FGF9*, *RUNX1*, *FOXL2*, *RSPO1* and *WNT4* in the different conditions was evaluated by RT-qPCR as described in Figure 2. Overall WT DHX37 is associated with increased levels of sex-determining transcripts in iSLCs compared with the mutant p.R674Q mutant.

### Changes in global gene expression following overexpression of DHX37 WT or mutant in iSLCs

We determined the transcriptomic signatures associated with a 48h over-expression of the WT or p.R674Q mutant proteins in iSLCs following transfection (**Figure 4A**). As shown in **Figure 4B**, the integration and projection of single-cell full-length RNA-seq data of the three samples on a common UMAP did not reveal any difference between the three conditions. Since iSLCs have the transcriptome signature of a homogeneous cell population, to gain statistical power, we aggregated their expression profiles using a pseudobulk approach with Seurat^42^ (AggregateExpression). The WT and mutant pseudobulk data were then compared. This resulted in the identification of 730 differentially expressed genes (DEGs), 341 up-regulated and 389 down-regulated in cells transfected with the mutant protein (**Figure 4C; Supp Table S1**). Gene enrichment analysis of the 730 DEGs indicated that amongst the ten most significant non-redundant biological processes were “Negative regulation of transcription, DNA-templated”, “RNA processing” and “Ribonucleoprotein complex biogenesis” (**Figure 4D**). These are consistent with the known biological functions of DHX37. The other significant non-redundant biological processes were “Protein transport”, “Cell death”, “Mitochondrion organization”, “Regulation of cell cycle”, “Regulation of translation”, “Protein targeting” and “Golgi vesicle transport”.

**Figure 4:**
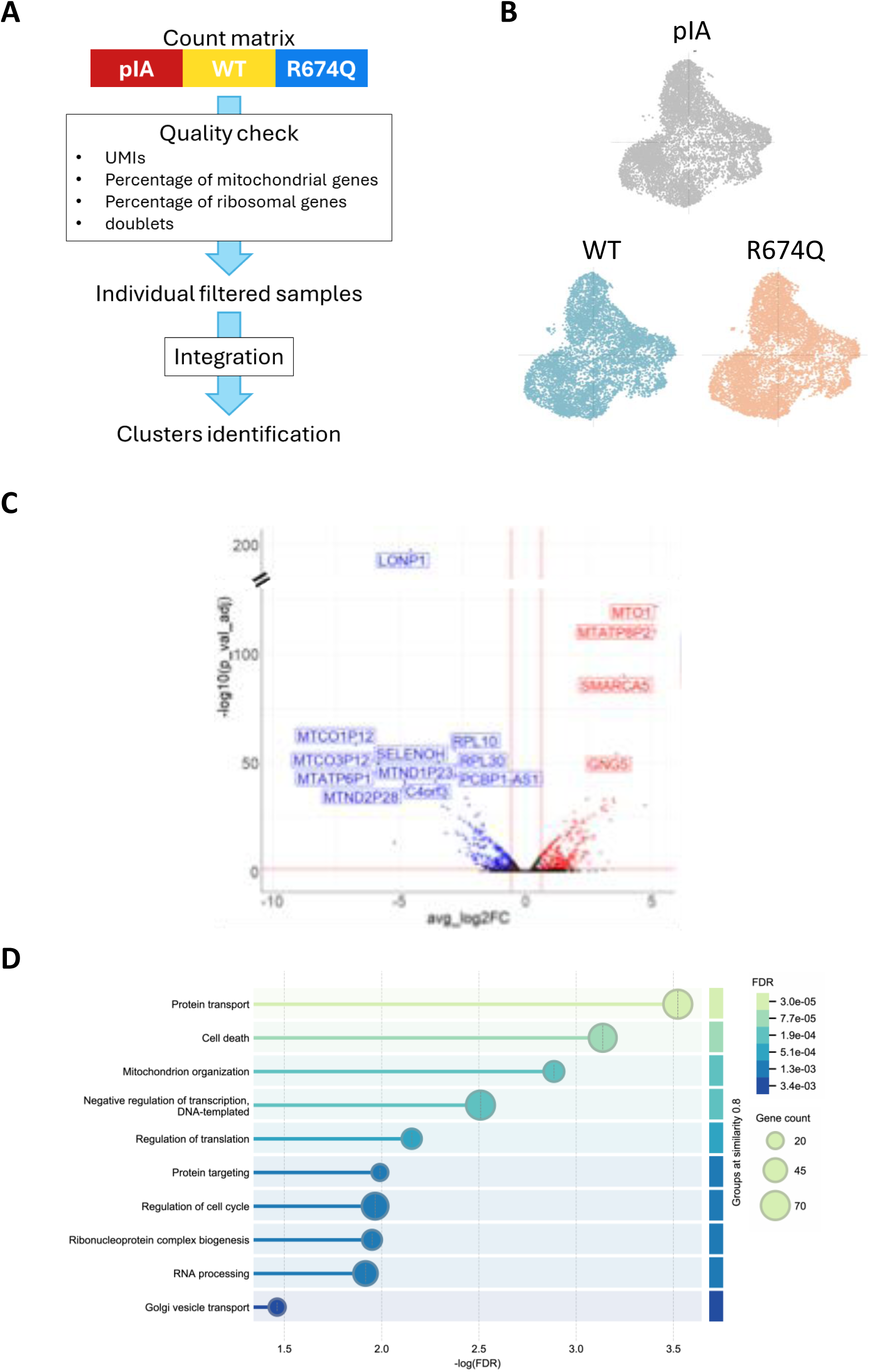
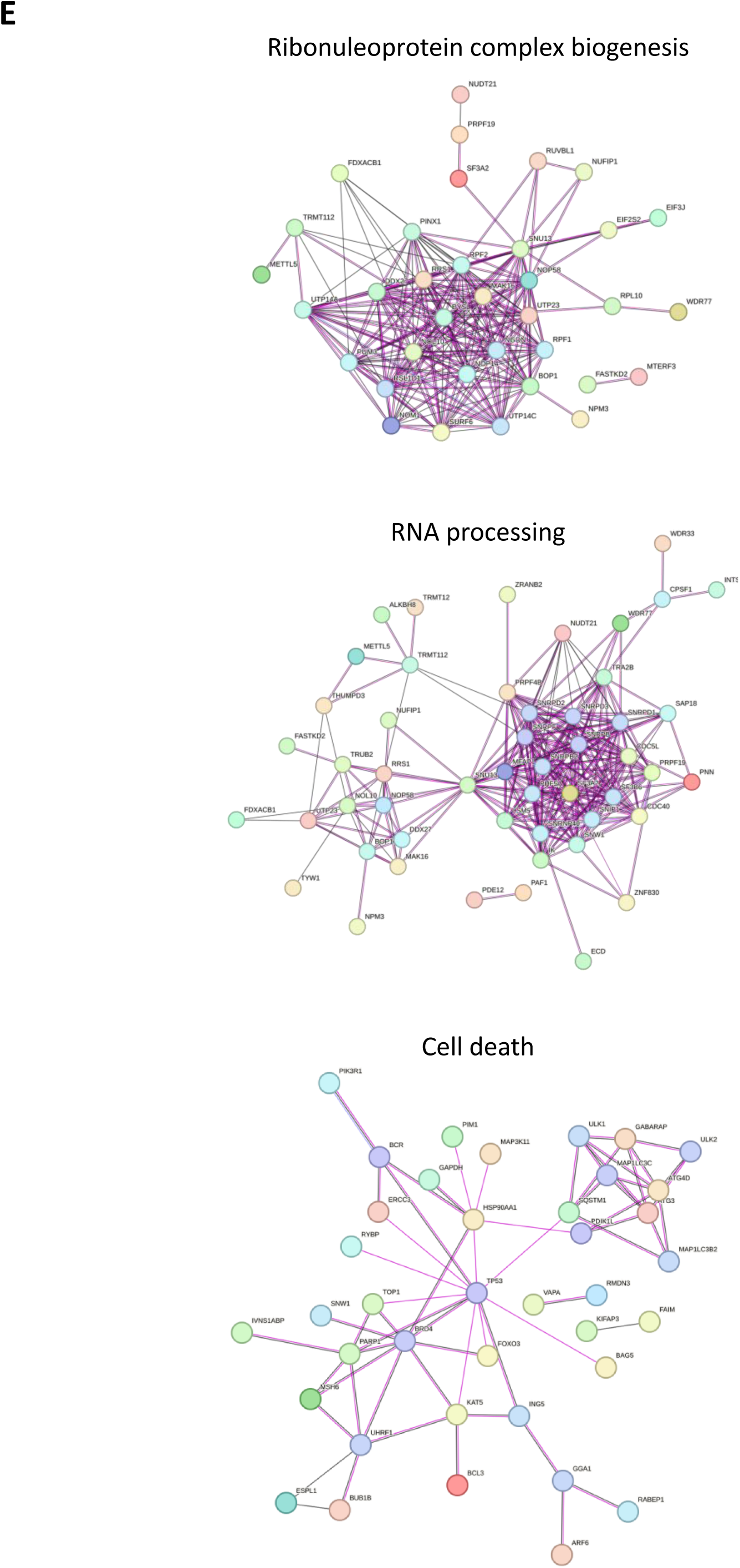
Comparison of WT and p.R674Q expression profiles confirmed known DHX37 functions and revealed dysregulation of genes implicated in new processes. **(A)** Workflow of global DEGs analysis. The three samples went through classical quality check and were then integrated. DEGs between clusters on the integrated UMAP were then defined. **(B**) UMAP showing clustering of the cells in the three conditions: pIA, WT and p.R674Q. Samples were homogeneous and no sample-specific cluster was observed. **(C)** Volcano plot showing the most significant altered gene expression profiles. Blue indicates the transcripts with upregulated expression profiles following transfection with the p.R674Q DHX37 construct in iSLCs, whereas red indicates the transcripts upregulated following transfection with the WT DHX37. **(D)** GO enrichment analysis of DEGs. GO enrichment analysis was performed using STRING. 602 out of 730 DE transcripts were considered (see **Supp Table S4**). The top 10 most significant non-redundant GO terms with less than 70 genes are indicated. **(E)** Gene networks based on 3 of the significant GO enrichments. The nodes in blue indicate an upregulation following transfection with the mutant DHX37, nodes in red indicate an upregulation following transfection with WT DHX37.

### WT DHX37 target transcripts identified by HyperTRIBE shows enrichment for cytoskeleton-based gene ontologies

DHX37 is widely expressed (https://www.proteinatlas.org/ENSG00000150990-DHX37), however its expression is low in human fetal Sertoli cells (https://www.reproductivecellatlas.org/gonads/human-main-male/)^48^. We found that *DHX37* expression increases during the course of XY iPSC differentiation towards iSLCs (**Supp Fig S3**). To define RNA transcripts physically interacting with the DHX37 protein, we used the HyperTRIBE (Targets of RNA-binding proteins Identified By Editing) approach. This protocol has proven to be powerful to identify the targets of RNA-binding proteins^49–51^. Here, we combined the use of HyperTRIBE and single-cell full-length RNA-seq, to enable an analysis of the entire transcript sequence for the identification of A>G SNPs. After validation of the HyperTRIBE fusion proteins and localization of the fusion protein in HEK-293T cells (**Supp Fig S1**), constructs were transfected into iSLCs. After 48h, transfected cells were sorted based on GFP expression (**Supp Fig S1C**) and after single-cell isolation of the cDNAs, the libraries were prepared for full-length RNA-sequencing. This approach required the development of specific bioinformatic pipelines (**Supp Figure S4A**). We obtained 7,507, 13,060 and 12,602 filtered variants (**Supp Table S2**) in 3,030, 4,892 and 5,139 transcripts in the pIA (negative control), WT and p.R674Q conditions respectively (**Supp Table 3**, **Supp Figure S4B**). As expected, the number of variants observed in the pIA was lower than in the other conditions as the ADAR2 catalytic site alone is not able to interact with transcripts. The variants observed in the pIA conditions may correspond to endogenous editing that naturally occurs in the cells^52^. We then focused the analysis on the transcripts having at least three A>G editing events (**Supp Figure S4C**).

We obtained 785 transcripts matching the filtering parameters in the WT condition (**Supp Table S3**). Pseudogenes, novel transcripts or long non-coding RNAs were filtered from the group to obtain a subset of 372 transcripts were used to performed a Gene Ontology (GO) enrichment analysis by Search Tool for the Retrieval of Interacting Genes/Proteins (STRING) (https://string-db.org/)^43^. We found significant enrichment for the biological processes of “regulation of cell motility”, “cell projection organization”, “regulation of plasma membrane organization” and “cytoskeleton organization” (**Figure 5A, Supp Table S4**). These data suggest that wild-type DHX37 interacts with transcripts involved in connected pathways, where cytoskeleton organization is central. Of the 372 edited transcripts analyzed, 148 are interacting within gene regulatory networks (**Figure 5B**). *JAK2*, *NUDCD1*, *POLR2A*, *PTK2*, *PSMA2* and *PSMB2* constituted the main nodes (more than 10 interactions) in this network. Of interest, PTK2 (Protein Tyrosine Kinase 2) is a tyrosine kinase localized at the focal adhesions that plays multiple roles including cell migration, adhesion, reorganization of the actin cytoskeleton and cell cycle progression^53^. NUDCD1 (NudC Domain Containing 1) is a tumor associated antigen with expression in normal tissues restricted to testis^54^ where its function has not been yet characterized.

**Figure 5:**
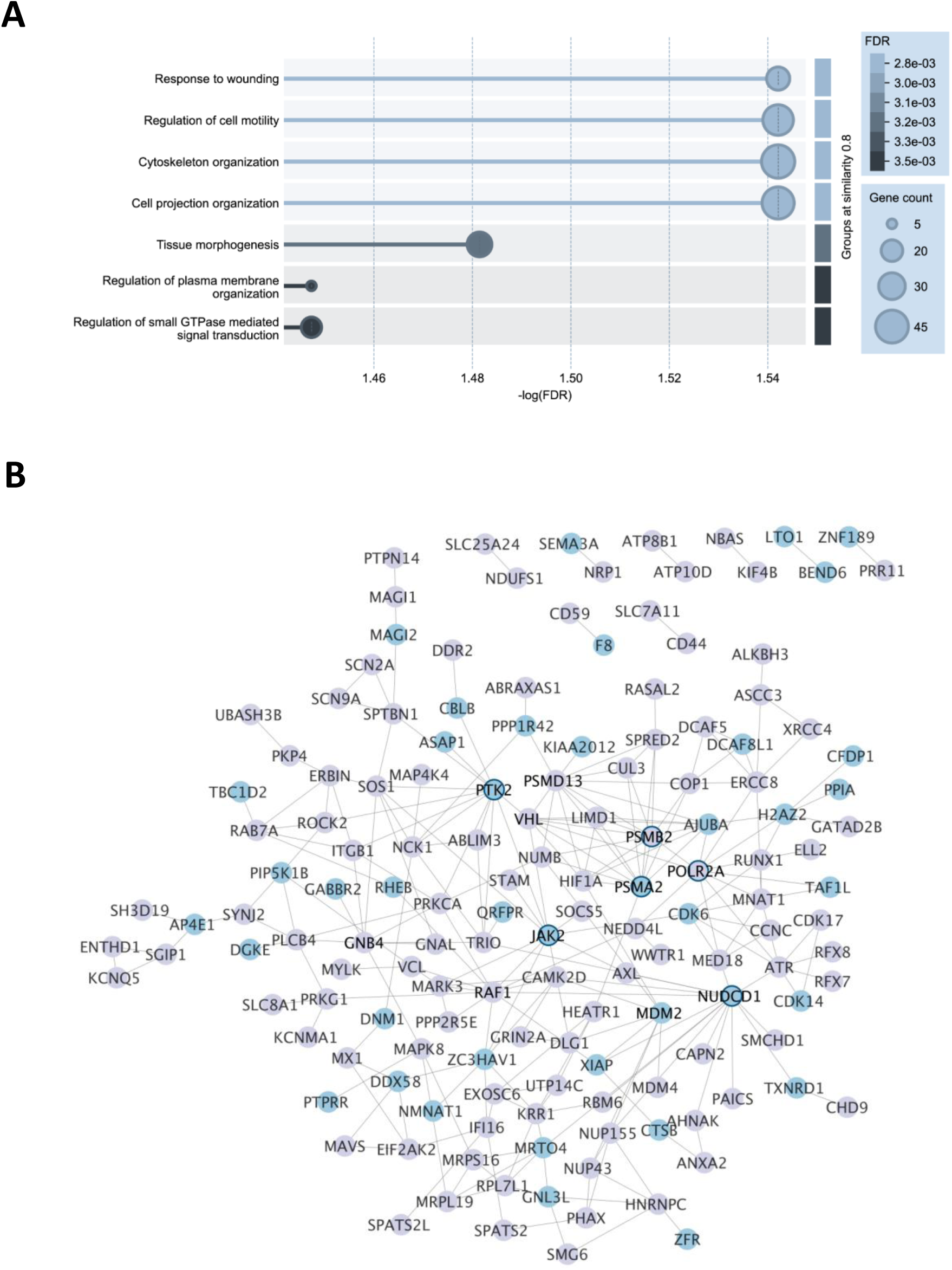
WT DHX37 target transcripts identified by HyperTRIBE are involved in cytoskeleton-based processes. **(A)** GO enrichment analysis of the 372 filtered transcripts edited in presence of WT DHX37 using STRING. The non-redundant GO terms are shown (see **Supp Table S4** for the complete list). **(B)** The transcripts edited in presence of WT DHX37 interact within a regulatory network. The network was built with STRING, considering “Experiments” and “Databases” as active interactive sources. The minimum required interaction score was set to medium and disconnected nodes were hidden. Formatting was performed using Cytoscape. Blue nodes indicate transcripts interacting specifically with the WT variant. Grey nodes were found to interact with both the WT and the p.R674Q mutant.

A comparison of the DEGs between the WT and DHX37 p.R674Q proteins (**Supp Table S5**) with the edited transcripts revealed 25 that were edited at least three times by the DHX37-ADAR2_cd_ construct, of which five (*LOXL2*, *LTO1*, *CBLB*, *KAT8* and *LRRC28*) were specifically interacting with WT DHX37.

### Specific targets of DHX37 p.R674Q mutant revealed by HyperTRIBE

Forty-eight hours following transfection of the p.R674Q construct in the iSCLs, transfected cells were sorted based on GFP expression and, after single-cell isolation of the cDNAs, the libraries were prepared for full-length RNA-seq. We observed 894 transcripts with at least 3 A>G SNPs, with 397 of these transcripts being specific to the mutant protein (**Supp Table S2**, **Figure 6A**). Six of them (*EPB41L2*, *KATNAL1*, *MARK4*, *NEDD4*, *NF1*, and *SIAH1*) are associated with testicular phenotypes in mice or humans (**Supp Table S6**).

**Figure 6:**
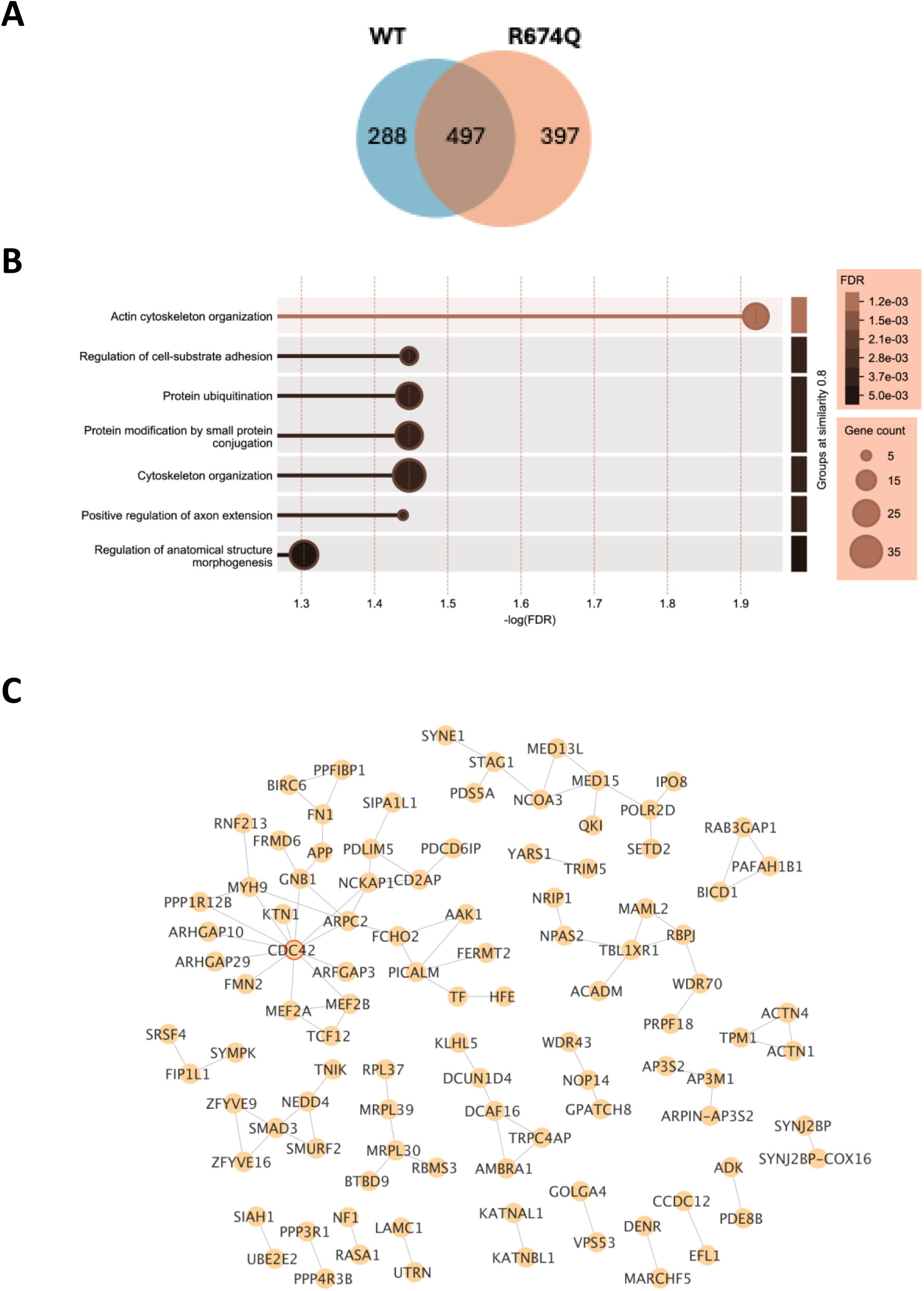
HyperTRIBE identification of specific targets of DHX37 p.R674Q mutant. **(A)** Venn diagram showing overlap between the transcripts interacting with WT DHX37 and p.R674Q DHX37. **(B)** GO enrichment analysis of the transcripts specifically edited in presence of p.R674Q DHX37, using STRING. 397 out of 894 transcripts were considered for the analysis. The non-redundant GO terms are shown (see **Supp Table S4** for the complete list). **(C)** The transcripts edited in presence of p.R674Q DHX37 interact within a regulatory network. The network was built with STRING and formatted using Cytoscape in Figure 4.

Following filtering of pseudogenes, novel transcripts or long non-coding RNAs, 266 edited transcripts were considered for the GO analysis. We observed a significant enrichment for the biological processes “actin cytoskeleton organization”, “cytoskeleton organization” or “regulation of cell-substrate adhesion” (**Figure 6B**). The GO terms “protein-ubiquitination” and “protein modification by small protein conjugation” were specific to the p.R674Q mutant. Of the 266 edited transcripts analyzed, 101 were interacting within gene regulatory networks (**Figure 6C**) where CDC42, a central protein in the establishment of cell polarity^55^ that is necessary for Sertoli cell survival in mice^56^, constituted the main node. Another major node was focused on ACTN4, an actin-binding protein linking F-actin to the membrane.

The comparison of the transcripts that interact with mutant DHX37 together with the DEGs (**Supp Table S5**) revealed nine genes in common (*C1orf21*, *CSAD*, *GALC*, *MB21D2*, *N4BP1*, *NCKAP1*, *UHMK1*, *ZNF626* and *ZNFX1*).

### Overexpression of WT or p.R674Q DHX37 in iSLCs did not impact selected targeted transcripts

The HyperTRIBE datasets of transcripts interacting with either the WT or p.R674Q DHX37 proteins indicated a bias for genes involved in the regulation of cytoskeleton-based processes. We sought to determine whether the interaction of DHX37 variants with selected transcripts altered their expression profiles (**Figure 7A**). We focused the analysis on *PTK2*, a transcript targeted by the WT protein, and *CDC42* and *ACTN4*, both targets of the mutated p.R674Q protein, because they constitute the main nodes in the gene networks regulated by DHX37. We observed that *PTK2* expression was increased after the introduction of both WT (fold change = 1.70) and R674Q DHX37 (fold change = 1.74) in iSLCs. Only WT DHX37 had a significant impact on *ACTN4* (fold change = 1.67) and none of the variants had any impact on *CDC42* (**Figure 7B**). Therefore, interaction with either the WT or mutant DHX37 proteins was not necessarily followed by changes in the expression profile of these genes after the 48h-transfection period.

**Figure 7:**
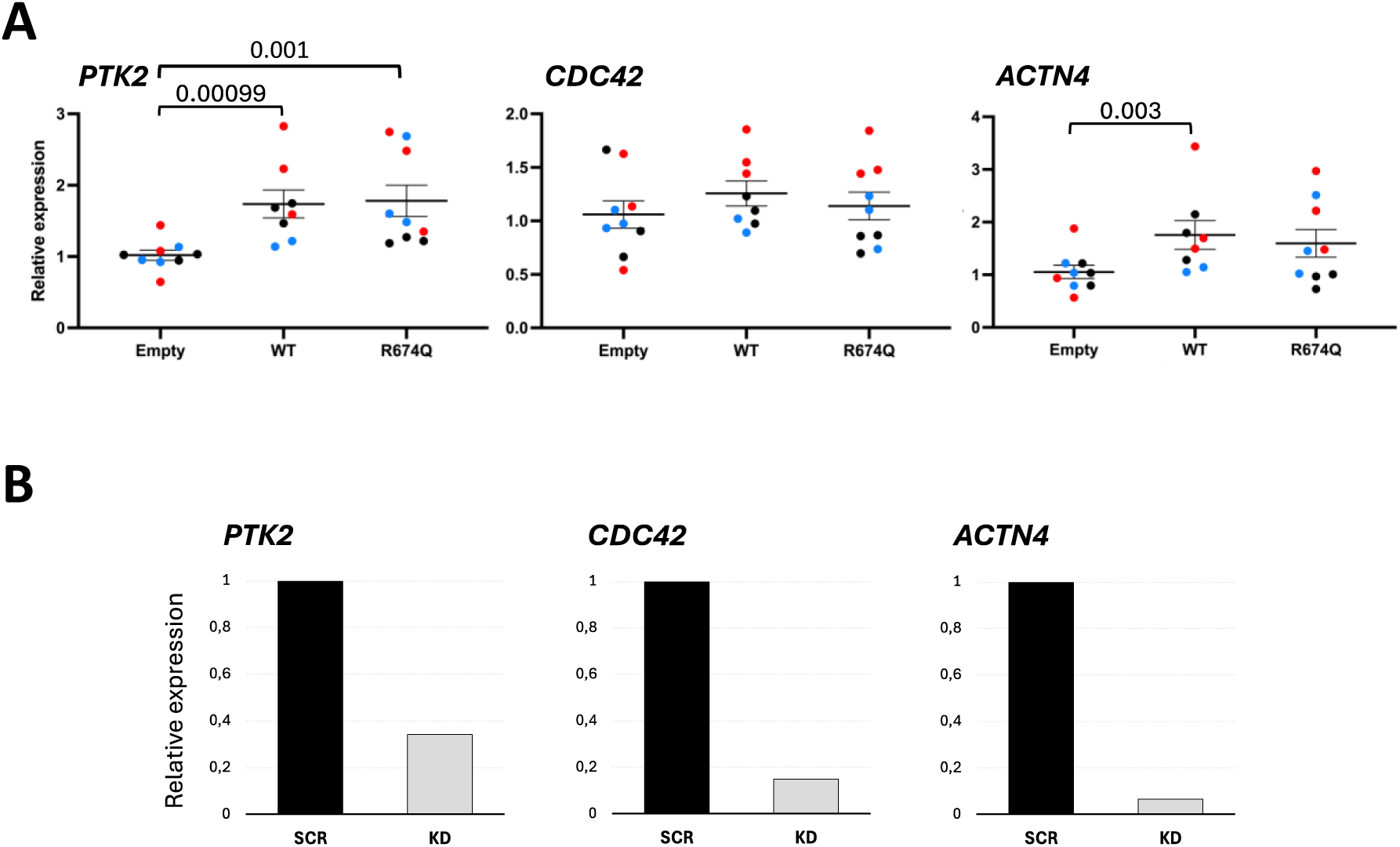
Dysregulation of DHX37 expression did not impact candidate targeted transcripts. **(A)** Effect of overexpression of DHX37 WT and p.R674Q variants on *PTK2*, *CDC42* and *ACTN4* in iSLCs. **(B)** Verification of *PTK2*, *CDC42* and *ACTN4* KD efficiencies by RT-qPCR. Levels of expression in the different conditions was evaluated by RT-qPCR as described In Figure 2.

## Discussion

DHX37 is an evolutionarily conserved RNA helicase, crucial for ribosome biogenesis from yeast to humans. Recent studies have revealed additional tissue specific functions for DHX37 that include regulation of splicing and transcription^31–33^. Variants in *DHX37* are a frequent cause of human male-to-female sex-reversal and testicular regression syndrome^19^ but the mechanism(s) by which they may engender these phenotypes is unknown. The variants associated with testicular phenotypes impact the RecA1 and RecA2 functional motifs of the protein^23^, however, the mutant proteins retain ATPase activity (**Figure 1**) suggesting that the mutations are highly unlikely to affect a core function of DHX37. This is corroborated by the patients’ phenotype where, other than the gonad, no other tissue is affected.

The overexpression of WT DHX37 and one of the most common recurrent mutations, p.R674Q, in iSLCs altered gonad specific gene expression. The WT DHX37 resulted in a significant increase in expression of genes involved in testes-determination and development. As compared to WT DHX37, overexpression of p.R674Q resulted in reduced expression of several pro-testis genes (*SOX9* and *AMH)* and an increase in expression of pro-ovary genes (*RUNX1*, *RSPO1* and *FOXL2)* (**Figure 3**). Murine and human studies have established that testis-determination occurs in a small number of supporting cells and requires a minimum threshold level of pro-testis gene expression to be reached within a critical, restricted developmental window. An *Sry* transgene model has established this critical window, that overlaps with a transient *Sry* gene expression, to a 6-h interval in mice (11.0 -11.25 dpc)^57^. A delay, either in reaching peak expression levels^58–61^, or in the initiation of expression (after 11.3 dpc)^57^ or premature upregulation of pro-ovarian genes before *Sry* expression results in XY sex-reversal^10^. A twofold or less reduction in murine *Sry* expression is sufficient to cause XY sex-reversal^62^. Studies of human XY male-to-female sex-reversal also show that gene expression threshold levels are critical for correct testis-determination^63,64^. The study of cryptic changes in the SRY protein, observed in rare familial cases of human sex-reversal that is transmitted by fertile fathers, have demonstrated that SRY function is exquisitely sensitive to critical threshold levels^64,64,65^. The commitment of the gonadal supporting cell lineage to Sertoli cells is also highly dependent on a threshold density of pre-Sertoli cells^66,67^. Therefore, the changes in expression profiles of pro-testis and pro-ovarian genes that we observed after 48 hours of transfection of either WT or p.R674Q DHX37 proteins in iSLCs may contribute to the phenotype of testicular dysgenesis in DHX37 patients.

Human DHX37 and its yeast ortholog Dhr1 are both RNA-helicases that physically interact with RNA species^26,27^. In zebrafish, Dhx37 interacts with glycine receptor mRNA transcripts^33^. We defined transcripts targeted by either the DHX37, WT or p.R674Q, in iSLCs using HyperTRIBE followed by single-cell full-length RNA-sequencing. In both conditions, we found that transcripts involved in cellular processes related to cytoskeleton organization or regulation were significantly enriched. The transcripts, *PTK2* (specific for WT DHX37) and *CDC42 and ACTN4* (both specific for the p.R674Q protein) are central in the network of transcripts interacting with either the WT or mutant RNA-helicase (**Figures 5 and 6**). These factors are important for adult murine Sertoli cells, regulating the blood-testis-barrier function or even cell survival^56,68–70^. These three transcripts are expressed in human fetal Sertoli cells (https://www.reproductivecellatlas.org/)^48^. Disruption of the biological function of these factors could be deleterious for Sertoli cell function and may explain the 46,XY DSD or TRS phenotype in DHX37 patients. Furthermore, the p.R674Q DHX37 protein, but not the WT DHX37 protein, was observed to interact with *EPB41L2*, *KATNAL1*, *MARK4*, *NEDD4*, *NF1*, and *SIAH1* mRNAs. Mouse knock-out models of these genes are associated with testicular phenotypes^71–82^ ranging from XY male-to-female sex-reversal to milder phenotypes of atypical male external genitalia or male infertility (**Supp Table S6**).

Although these results suggest possible mechanisms leading to male-to-female sex-reversal associated with DHX37 variants, other potential mechanisms that could explain the gonadal phenotype are possible. DHX37 has the capacity to regulate mRNA splicing in zebrafish^33^, although this has not been investigated in mammalian cells. An analysis of differential splicing events in iSLCs between the WT and mutant DHX37 proteins may be informative. This is particularly relevant since the DHX37 protein has been shown to physically interact with SART3 (Squamous cell carcinoma antigen recognized by T cells 3) in hepatocellular carcinoma cells^32^. SART3 is an RNA binding protein required for spliceosome function by recycling small nuclear RNAs during pre-mRNA splicing^83^. Bi-allelic variants in human *SART3* are associated with a complex phenotype, termed INDYGON syndrome, which includes 46,XY gonadal dysgenesis and neuronal anomalies^84^. The mechanism responsible for the male-to-female sex-reversal observed in patients carrying pathogenic SART3 variants is unknown since *ex-vivo* studies did not reveal a major disruption of splicing events (e.g. no increase in intron retention or exon skipping)^84^.

In conclusion, the data presented here provide evidence of multiple mechanisms that can explain how specific variants in DHX37 result in 46,XY sex-reversal. Whilst, these mutant proteins retain ATPase activity and are not associated with the distinctive signatures of nucleolar stress, they are associated with global changes in gene expression in iSLCs and with changes in interactions with specific RNA transcripts that are known to be required for the formation and maintenance of the supporting cell lineages of the human testis.

## Acknowledgements

We thank Valentina Libri (former head of the Single Cell platform at Institut Pasteur) for her valuable advice regarding single-cell technology possibilities when the project started, as well as Chloé Mayère (University of Geneva) and Almira Chervova (Institut Pasteur) for advices regarding the computational analyses.

## Author contributions

M.E., A.B. and K.M. conceived the experiments. M.E., L.S., S.W., J.B.-T., C.E., C.B. and V.S. performed the experiments. M.E., E.T., and E.K. analyzed the RNA-seq data and performed the computational analyzes. P.-H.C. conducted the flow cytometric analysis. M.E. collated the experimental data. The manuscript was written by M.E., E.T., K.M. and A.B. All the authors read and agreed with the data being presented in the manuscript.

## Conflicts of interest

The authors declare that they have no competing interests.

## Data availability

All data needed to evaluate the conclusions in the article are included in the paper and/or the Supplementary Information. The single-cell full-length RNA-seq raw files (fastq) dataset is being deposited to the Gene Expression Omnibus (GEO) repository.

## Funding

M.E., A.B. and K.M are funded by the Agence Nationale de la Recherche (ANR), ANR-24-CE13-4205-01, ANR-10-LABX-73 REVIVE, ANR-19-CE14-0012, ANR-23-CE14-0061, and ANR-23-CE14-0068. In the interest of open-access publication, the authors apply a CC-BY open-access license to any manuscript accepted for publication (AAM) resulting from this submission.

## Supplementary Tables and Figure legends

**Supp Table S1: Differentially expressed genes between iSLCs expressing WT DHX37 and p.R674Q DHX37**. Lists of (i) all DEGs and (ii) the DEGs considered by STRING for GO enrichment analysis are provided.

**Supp Table S2: A>G SNPs positions in WT and p.R674Q DHX37 conditions**. The positions of SNPs in transcripts, which have at least three editing events in at least one of the two conditions are indicated.

**Supp Table S3: Lists of edited transcripts.** (i) the list of all the transcripts with A>G SNPs in WT and p.R674Q conditions, (ii) the list of transcripts with minimum of 3 A>G SNPs in WT or p.R674Q conditions, (iii) the list of 372 transcripts interacting with WT DHX37, retained by STRING for GO enrichment analysis, and (iv) the list of 397 transcripts specifically interacting with p.R674Q DHX37, retained by STRING for GO enrichment analysis.

**Supp Table S4: GO enrichment analyses**. The significant GO terms involving (i) the transcripts interacting with WT DHX37, (ii) the transcripts interacting specifically with p.R674Q DHX37, and (iii) the DEGs in iSLCs after introduction of WT or p.R674Q DHX37 are listed. Only the non-redundant GO terms comprising less than 70 genes were displayed in the Figures, to avoid generic terms.

**Supp Table S5: Overlap between DE transcripts and transcripts targeted by DHX37 variants**. The list indicates all the overlapping transcripts, as well as the condition they were found edited (WT, p.R674Q or both).

**Supp Table S6: Review of testicular phenotypes associated with *EPB41L2*, *KATNAL1*, *MARK4*, *NEDD4*, *NF1*, *PAFAH1B1* and *SIAH1***. The table indicates the murine or human testicular phenotypes associated with these genes in the literature.

---

**Supp. Fig. S1.**
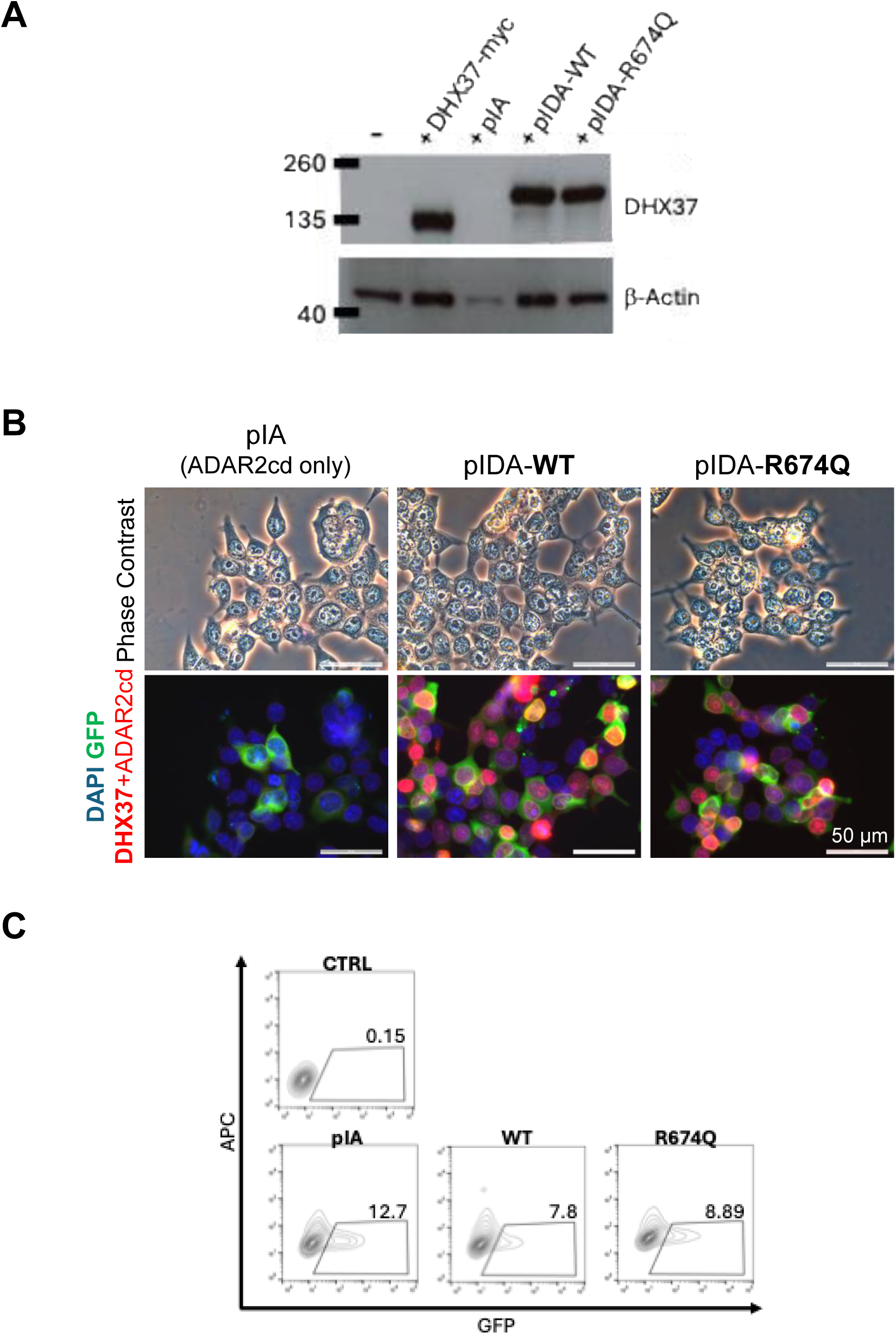

**Supp. Fig. S2.**
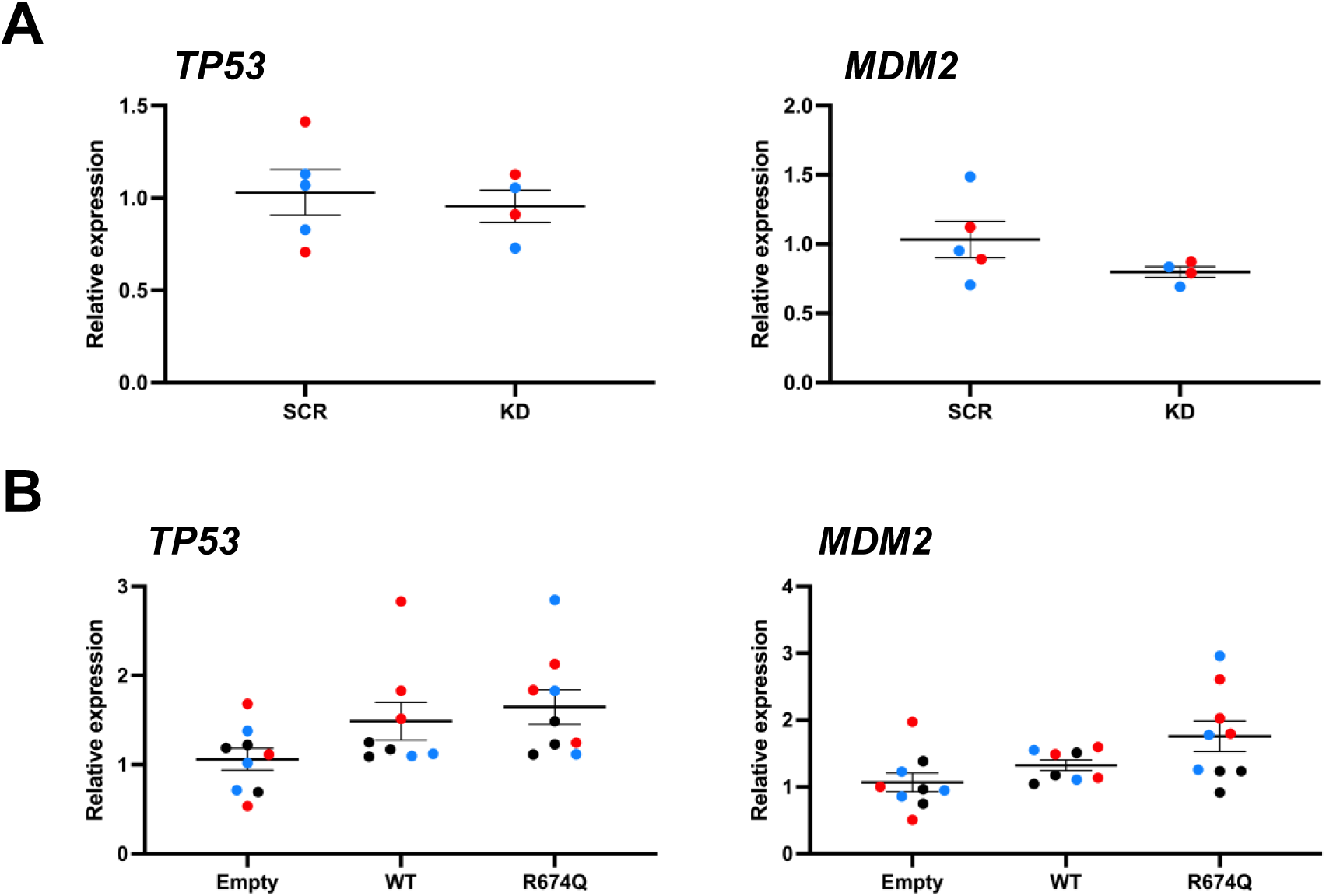

**Supp. Fig. S3.**
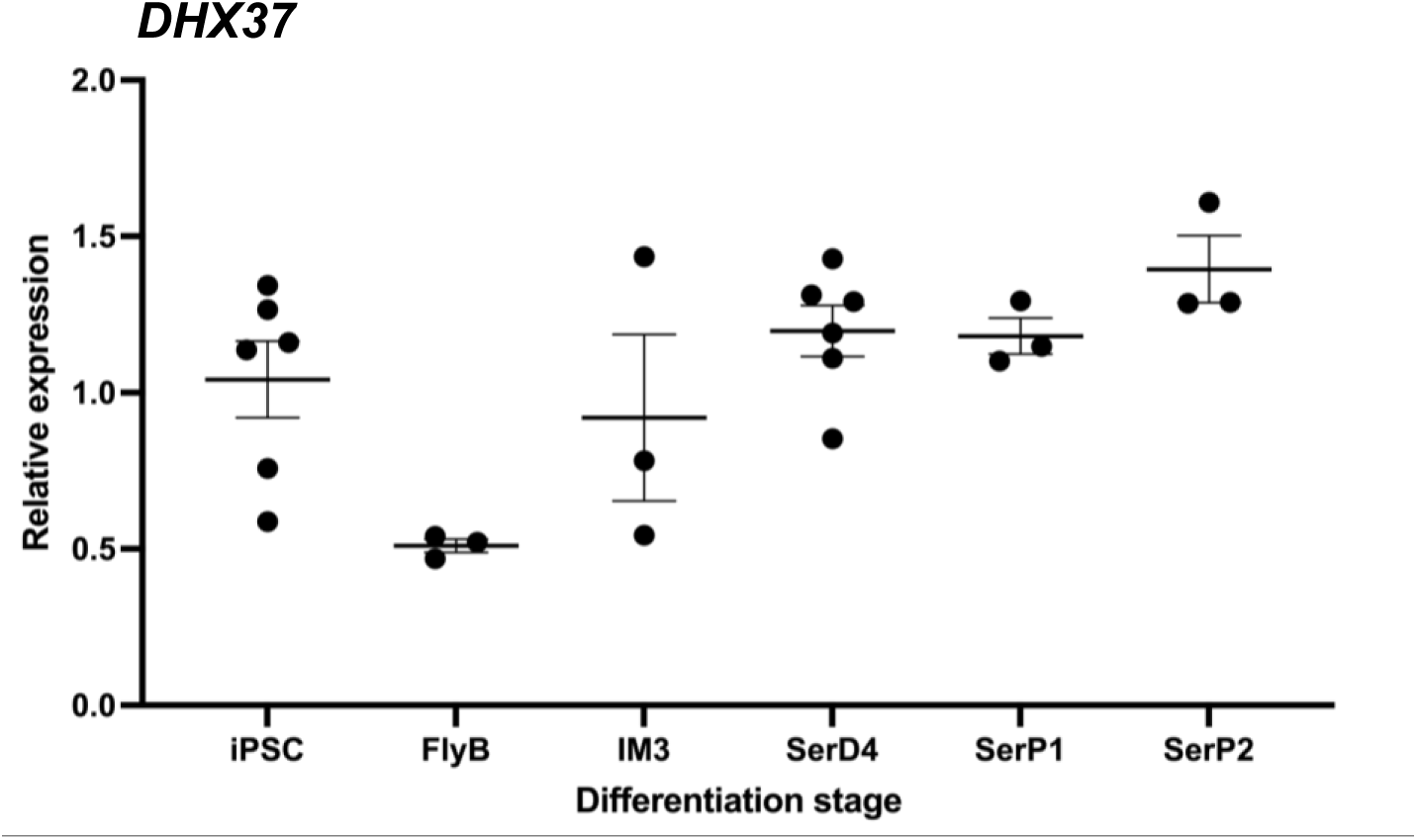

**Supp. Fig. S4.**
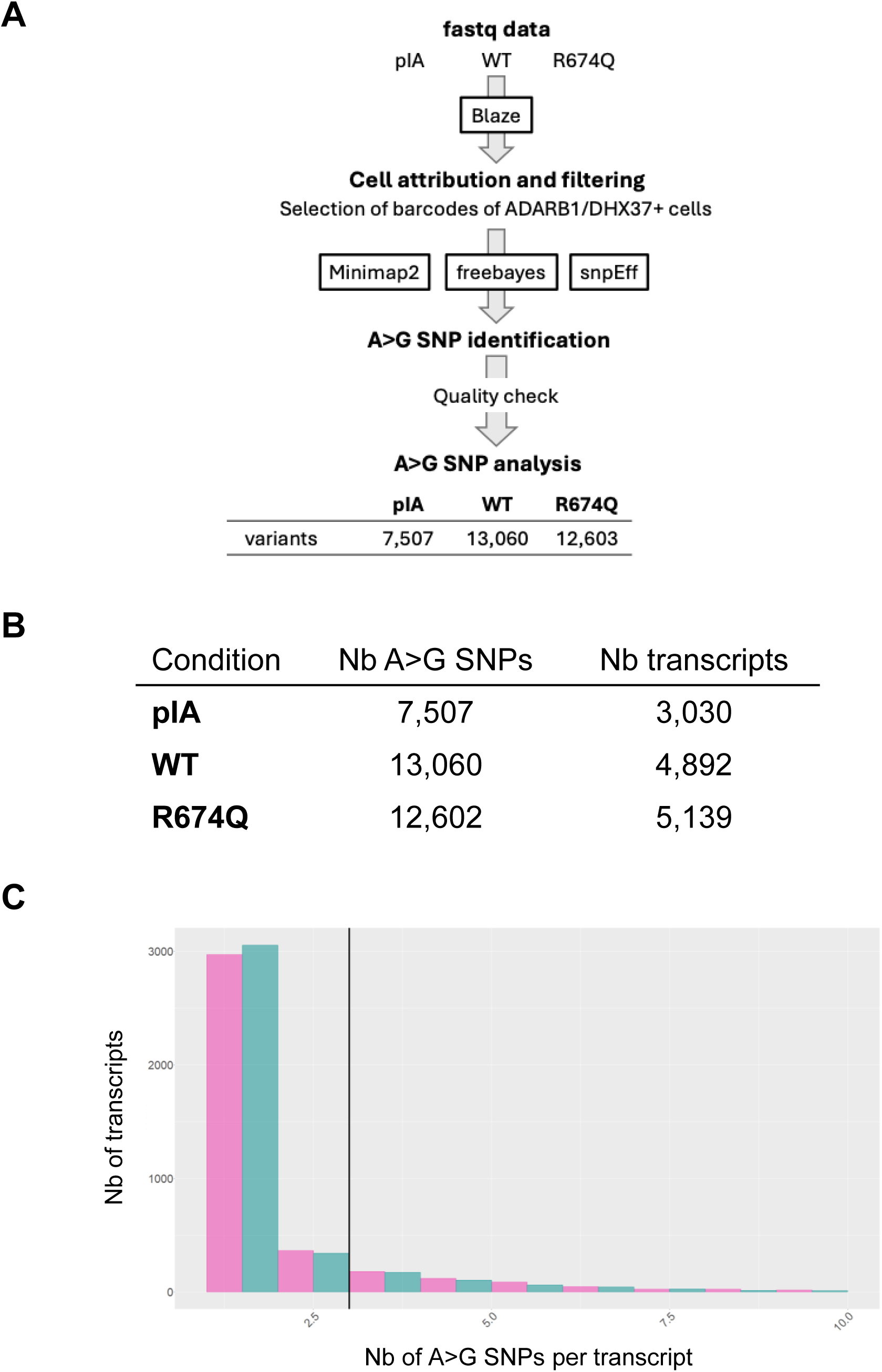

## Notes

### Competing Interest Statement

The authors have declared no competing interest.

## References

1 Capel B. Vertebrate sex determination: evolutionary plasticity of a fundamental switch. Nat Rev Genet 2017;18:675–89. 10.1038/nrg.2017.60.

2 Sekido R, Bar I, Narváez V, Penny G, Lovell-Badge R. SOX9 is up-regulated by the transient expression of SRY specifically in Sertoli cell precursors. Dev Biol 2004;274:271–9. 10.1016/j.ydbio.2004.07.011.

3 Kim Y, Kobayashi A, Sekido R, DiNapoli L, Brennan J, Chaboissier M-C, et al. Fgf9 and Wnt4 act as antagonistic signals to regulate mammalian sex determination. PLoS Biol 2006;4:e187. 10.1371/journal.pbio.0040187.

4 Kim Y, Bingham N, Sekido R, Parker KL, Lovell-Badge R, Capel B. Fibroblast growth factor receptor 2 regulates proliferation and Sertoli differentiation during male sex determination. Proc Natl Acad Sci U S A 2007;104:16558–63. 10.1073/pnas.0702581104.

5 Bagheri-Fam S, Sim H, Bernard P, Jayakody I, Taketo MM, Scherer G, et al. Loss of Fgfr2 leads to partial XY sex reversal. Dev Biol 2008;314:71–83. 10.1016/j.ydbio.2007.11.010.

6 Wilhelm D, Martinson F, Bradford S, Wilson MJ, Combes AN, Beverdam A, et al. Sertoli cell differentiation is induced both cell-autonomously and through prostaglandin signaling during mammalian sex determination. Dev Biol 2005;287:111–24. 10.1016/j.ydbio.2005.08.039.

7 Malki S, Nef S, Notarnicola C, Thevenet L, Gasca S, Méjean C, et al. Prostaglandin D2 induces nuclear import of the sex-determining factor SOX9 via its cAMP-PKA phosphorylation. EMBO J 2005;24:1798–809. 10.1038/sj.emboj.7600660.

8 Moniot B, Declosmenil F, Barrionuevo F, Scherer G, Aritake K, Malki S, et al. The PGD2 pathway, independently of FGF9, amplifies SOX9 activity in Sertoli cells during male sexual differentiation. Development 2009;136:1813–21. 10.1242/dev.032631.

9 Nel-Themaat L, Jang C-W, Stewart MD, Akiyama H, Viger RS, Behringer RR. Sertoli cell behaviors in developing testis cords and postnatal seminiferous tubules of the mouse. Biol Reprod 2011;84:342–50. 10.1095/biolreprod.110.086900.

10 Gregoire EP, De Cian M-C, Migale R, Perea-Gomez A, Schaub S, Bellido-Carreras N, et al. The -KTS splice variant of WT1 is essential for ovarian determination in mice. Science 2023;382:600–6. 10.1126/science.add8831.

11 Nicol B, Grimm SA, Chalmel F, Lecluze E, Pannetier M, Pailhoux E, et al. RUNX1 maintains the identity of the fetal ovary through an interplay with FOXL2. Nat Commun 2019;10:5116. 10.1038/s41467-019-13060-1.

12 Boulanger L, Pannetier M, Gall L, Allais-Bonnet A, Elzaiat M, Le Bourhis D, et al. FOXL2 is a female sex-determining gene in the goat. Curr Biol 2014;24:404–8. 10.1016/j.cub.2013.12.039.

13 Vainio S, Heikkilä M, Kispert A, Chin N, McMahon AP. Female development in mammals is regulated by Wnt-4 signalling. Nature 1999;397:405–9. 10.1038/17068.

14 Chassot A-A, Ranc F, Gregoire EP, Roepers-Gajadien HL, Taketo MM, Camerino G, et al. Activation of beta-catenin signaling by Rspo1 controls differentiation of the mammalian ovary. Hum Mol Genet 2008;17:1264–77. 10.1093/hmg/ddn016.

15 Maatouk DM, DiNapoli L, Alvers A, Parker KL, Taketo MM, Capel B. Stabilization of beta-catenin in XY gonads causes male-to-female sex-reversal. Hum Mol Genet 2008;17:2949–55. 10.1093/hmg/ddn193.

16 Hughes IA, Houk C, Ahmed SF, Lee PA, Group LC. Consensus statement on management of intersex disorders. Arch Dis Child 2006;91:554–63. 10.1136/adc.2006.098319.

17 Josso N, Briard ML. Embryonic testicular regression syndrome: variable phenotypic expression in siblings. J Pediatr 1980;97:200–4. 10.1016/s0022-3476(80)80474-4.

18 Marcantonio SM, Fechner PY, Migeon CJ, Perlman EJ, Berkovitz GD. Embryonic testicular regression sequence: a part of the clinical spectrum of 46,XY gonadal dysgenesis. Am J Med Genet 1994;49:1–5. 10.1002/ajmg.1320490102.

19 McElreavey K, Jorgensen A, Eozenou C, Merel T, Bignon-Topalovic J, Tan DS, et al. Pathogenic variants in the DEAH-box RNA helicase DHX37 are a frequent cause of 46,XY gonadal dysgenesis and 46,XY testicular regression syndrome. Genet Med 2020;22:150–9. 10.1038/s41436-019-0606-y.

20 Zidoune H, Martinerie L, Tan DS, Askari M, Rezgoune D, Ladjouze A, et al. Expanding DSD Phenotypes Associated with Variants in the DEAH-Box RNA Helicase DHX37. Sex Dev 2021;15:244–52. 10.1159/000515924.

21 Shimura K, Ichihashi Y, Nakano S, Sato T, Hamajima T, Numasawa K, et al. DHX37 Variant is One of Common Genetic Causes in Japanese Patients with Testicular Regression Syndrome / Partial Gonadal Dysgenesis without Müllerian Derivatives. Horm Res Paediatr 2024. 10.1159/000537761.

22 Yang H, Ma X, Tian H, Yuan J, Wu D, Dong G, et al. Two Novel Heterozygous Variants in RecA2 Domain of DHX37 Cause 46,XY Gonadal Dysgenesis and Testicular Regression Syndrome. Sex Dev 2023;17:198–202. 10.1159/000534086.

23 McElreavey K, Pailhoux E, Bashamboo A. DHX37 and 46,XY DSD: A New Ribosomopathy? Sex Dev 2022;16:194–206. 10.1159/000522004.

24 Sloan KE, Bohnsack MT. Unravelling the Mechanisms of RNA Helicase Regulation. Trends Biochem Sci 2018;43:237–50. 10.1016/j.tibs.2018.02.001.

25 Karaca E, Harel T, Pehlivan D, Jhangiani SN, Gambin T, Coban Akdemir Z, et al. Genes that Affect Brain Structure and Function Identified by Rare Variant Analyses of Mendelian Neurologic Disease. Neuron 2015;88:499–513. 10.1016/j.neuron.2015.09.048.

26 Sardana R, Liu X, Granneman S, Zhu J, Gill M, Papoulas O, et al. The DEAH-box helicase Dhr1 dissociates U3 from the pre-rRNA to promote formation of the central pseudoknot. PLoS Biol 2015;13:e1002083. 10.1371/journal.pbio.1002083.

27 Choudhury P, Hackert P, Memet I, Sloan KE, Bohnsack MT. The human RNA helicase DHX37 is required for release of the U3 snoRNP from pre-ribosomal particles. RNA Biol 2019;16:54–68. 10.1080/15476286.2018.1556149.

28 Roychowdhury A, Joret C, Bourgeois G, Heurgué-Hamard V, Lafontaine DLJ, Graille M. The DEAH-box RNA helicase Dhr1 contains a remarkable carboxyl terminal domain essential for small ribosomal subunit biogenesis. Nucleic Acids Res 2019;47:7548–63. 10.1093/nar/gkz529.

29 Phipps KR, Charette JM, Baserga SJ. The small subunit processome in ribosome biogenesis—progress and prospects. Wiley Interdiscip Rev RNA 2011;2:1–21. 10.1002/wrna.57.

30 Vanden Broeck A, Klinge S. An emerging mechanism for the maturation of the Small Subunit Processome. Curr Opin Struct Biol 2022;73:102331. 10.1016/j.sbi.2022.102331.

31 Dong MB, Wang G, Chow RD, Ye L, Zhu L, Dai X, et al. Systematic Immunotherapy Target Discovery Using Genome-Scale In Vivo CRISPR Screens in CD8 T Cells. Cell 2019;178:1189–1204.e23. 10.1016/j.cell.2019.07.044.

32 Liu Z, Ye Y, Liu Y, Liu Y, Chen H, Shen M, et al. RNA Helicase DHX37 Facilitates Liver Cancer Progression by Cooperating with PLRG1 to Drive Superenhancer-Mediated Transcription of Cyclin D1. Cancer Res 2022;82:1937–52. 10.1158/0008-5472.CAN-21-3038.

33 Hirata H, Ogino K, Yamada K, Leacock S, Harvey RJ. Defective escape behavior in DEAH-box RNA helicase mutants improved by restoring glycine receptor expression. J Neurosci 2013;33:14638–44. 10.1523/JNEUROSCI.1157-13.2013.

34 Gonen N, Eozenou C, Mitter R, Elzaiat M, Stévant I, Aviram R, et al. In vitro cellular reprogramming to model gonad development and its disorders. Sci Adv 2023;9:eabn9793. 10.1126/sciadv.abn9793.

35 Lebrigand K, Magnone V, Barbry P, Waldmann R. High throughput error corrected Nanopore single cell transcriptome sequencing. Nat Commun 2020;11:4025. 10.1038/s41467-020-17800-6.

36 Xu W, Rahman R, Rosbash M. Mechanistic implications of enhanced editing by a HyperTRIBE RNA-binding protein. RNA 2018;24:173–82. 10.1261/rna.064691.117.

37 You Y, Prawer YDJ, De Paoli-Iseppi R, Hunt CPJ, Parish CL, Shim H, et al. Identification of cell barcodes from long-read single-cell RNA-seq with BLAZE. Genome Biology 2023;24:66. 10.1186/s13059-023-02907-y.

38 Tian L, Jabbari JS, Thijssen R, Gouil Q, Amarasinghe SL, Voogd O, et al. Comprehensive characterization of single-cell full-length isoforms in human and mouse with long-read sequencing. Genome Biology 2021;22:310. 10.1186/s13059-021-02525-6.

39 Li H. Minimap2: pairwise alignment for nucleotide sequences. Bioinformatics 2018;34:3094–100. 10.1093/bioinformatics/bty191.

40 Garrison E, Marth G. Haplotype-based variant detection from short-read sequencing 2012. 10.48550/arXiv.1207.3907.

41 Cingolani P, Platts A, Wang LL, Coon M, Nguyen T, Wang L, et al. A program for annotating and predicting the effects of single nucleotide polymorphisms, SnpEff: SNPs in the genome of Drosophila melanogaster strain w1118; iso-2; iso-3. Fly 2012;6:80–92. 10.4161/fly.19695.

42 Hao Y, Stuart T, Kowalski MH, Choudhary S, Hoffman P, Hartman A, et al. Dictionary learning for integrative, multimodal and scalable single-cell analysis. Nat Biotechnol 2024;42:293–304. 10.1038/s41587-023-01767-y.

43 Szklarczyk D, Kirsch R, Koutrouli M, Nastou K, Mehryary F, Hachilif R, et al. The STRING database in 2023: protein-protein association networks and functional enrichment analyses for any sequenced genome of interest. Nucleic Acids Res 2023;51:D638–46. 10.1093/nar/gkac1000.

44 Shannon P, Markiel A, Ozier O, Baliga NS, Wang JT, Ramage D, et al. Cytoscape: a software environment for integrated models of biomolecular interaction networks. Genome Res 2003;13:2498–504. 10.1101/gr.1239303.

45 Schindelin J, Arganda-Carreras I, Frise E, Kaynig V, Longair M, Pietzsch T, et al. Fiji: an open-source platform for biological-image analysis. Nat Methods 2012;9:676–82. 10.1038/nmeth.2019.

46 Pfister AS, Kühl M. Of Wnts and Ribosomes. Prog Mol Biol Transl Sci 2018;153:131–55. 10.1016/bs.pmbts.2017.11.006.

47 Dannheisig DP, Bächle J, Tasic J, Keil M, Pfister AS. The Wnt/β-Catenin Pathway is Activated as a Novel Nucleolar Stress Response. J Mol Biol 2021;433:166719. 10.1016/j.jmb.2020.11.018.

48 Garcia-Alonso L, Lorenzi V, Mazzeo CI, Alves-Lopes JP, Roberts K, Sancho-Serra C, et al. Single-cell roadmap of human gonadal development. Nature 2022;607:540–7. 10.1038/s41586-022-04918-4.

49 Rahman R, Xu W, Jin H, Rosbash M. Identification of RNA-binding protein targets with HyperTRIBE. Nat Protoc 2018;13:1829–49. 10.1038/s41596-018-0020-y.

50 Nguyen DTT, Lu Y, Chu EL, Yang X, Park S-M, Choo Z-N, et al. HyperTRIBE uncovers increased MUSASHI-2 RNA binding activity and differential regulation in leukemic stem cells. Nat Commun 2020;11:2026. 10.1038/s41467-020-15814-8.

51 Piao W, Li C, Sun P, Yang M, Ding Y, Song W, et al. Identification of RNA-Binding Protein Targets with HyperTRIBE in Saccharomyces cerevisiae. Int J Mol Sci 2023;24:9033. 10.3390/ijms24109033.

52 Bazak L, Haviv A, Barak M, Jacob-Hirsch J, Deng P, Zhang R, et al. A-to-I RNA editing occurs at over a hundred million genomic sites, located in a majority of human genes. Genome Res 2014;24:365–76. 10.1101/gr.164749.113.

53 Le Coq J, Acebrón I, Rodrigo Martin B, López Navajas P, Lietha D. New insights into FAK structure and function in focal adhesions. J Cell Sci 2022;135:jcs259089. 10.1242/jcs.259089.

54 Yang XF, Wu CJ, McLaughlin S, Chillemi A, Wang KS, Canning C, et al. CML66, a broadly immunogenic tumor antigen, elicits a humoral immune response associated with remission of chronic myelogenous leukemia. Proc Natl Acad Sci U S A 2001;98:7492–7. 10.1073/pnas.131590998.

55 Etienne-Manneville S. Cdc42--the centre of polarity. J Cell Sci 2004;117:1291–300. 10.1242/jcs.01115.

56 Heinrich A, Bhandary B, Potter SJ, Ratner N, DeFalco T. Cdc42 activity in Sertoli cells is essential for maintenance of spermatogenesis. Cell Rep 2021;37:109885. 10.1016/j.celrep.2021.109885.

57 Hiramatsu R, Matoba S, Kanai-Azuma M, Tsunekawa N, Katoh-Fukui Y, Kurohmaru M, et al. A critical time window of Sry action in gonadal sex determination in mice. Development 2009;136:129–38. 10.1242/dev.029587.

58 Warr N, Carre G-A, Siggers P, Faleato JV, Brixey R, Pope M, et al. Gadd45γ and Map3k4 interactions regulate mouse testis determination via p38 MAPK-mediated control of Sry expression. Dev Cell 2012;23:1020–31. 10.1016/j.devcel.2012.09.016.

59 Warr N, Siggers P, May J, Chalon N, Pope M, Wells S, et al. Gadd45g is required for timely Sry expression independently of RSPO1 activity. Reproduction 2022;163:333–40. 10.1530/REP-21-0443.

60 Warr N, Siggers P, Carré G-A, Wells S, Greenfield A. Genetic Analyses Reveal Functions for MAP2K3 and MAP2K6 in Mouse Testis Determination. Biol Reprod 2016;94:103. 10.1095/biolreprod.115.138057.

61 Bullejos M, Koopman P. Delayed Sry and Sox9 expression in developing mouse gonads underlies B6-Y(DOM) sex reversal. Dev Biol 2005;278:473–81. 10.1016/j.ydbio.2004.11.030.

62 Albrecht KH, Young M, Washburn LL, Eicher EM. Sry expression level and protein isoform differences play a role in abnormal testis development in C57BL/6J mice carrying certain Sry alleles. Genetics 2003;164:277–88. 10.1093/genetics/164.1.277.

63 Elzaiat M, McElreavey K, Bashamboo A. Genetics of 46,XY gonadal dysgenesis. Best Pract Res Clin Endocrinol Metab 2022;36:101633. 10.1016/j.beem.2022.101633.

64 Chen Y-S, Racca JD, Weiss MA. Tenuous Transcriptional Threshold of Human Sex Determination. I. SRY and Swyer Syndrome at the Edge of Ambiguity. Front Endocrinol (Lausanne) 2022;13:945030. 10.3389/fendo.2022.945030.

65 Racca JD, Chatterjee D, Chen Y-S, Rai RK, Yang Y, Georgiadis MM, et al. Tenuous transcriptional threshold of human sex determination. II. SRY exploits water-mediated clamp at the edge of ambiguity. Front Endocrinol (Lausanne) 2022;13:1029177. 10.3389/fendo.2022.1029177.

66 Burgoyne PS, Buehr M, McLaren A. XY follicle cells in ovaries of XX XY female mouse chimaeras. Development 1988;104:683–8. 10.1242/dev.104.4.683.

67 Bunce C, Barske L, Zhang G, Capel B. Biased precursor ingression underlies the center-to-pole pattern of male sex determination in mouse. Development 2023;150:dev201060. 10.1242/dev.201060.

68 Wen Q, Tang EI, Gao Y, Jesus TT, Chu DS, Lee WM, et al. Signaling pathways regulating blood-tissue barriers - Lesson from the testis. Biochim Biophys Acta Biomembr 2018;1860:141–53. 10.1016/j.bbamem.2017.04.020.

69 Mori Y, Takashima S, Kanatsu-Shinohara M, Yi Z, Shinohara T. Cdc42 is required for male germline niche development in mice. Cell Rep 2021;36:109550. 10.1016/j.celrep.2021.109550.

70 Liu C, Wang H, Shang Y, Liu W, Song Z, Zhao H, et al. Autophagy is required for ectoplasmic specialization assembly in sertoli cells. Autophagy 2016;12:814–32. 10.1080/15548627.2016.1159377.

71 Taylor-Harris PM, Felkin LE, Birks EJ, Franklin RCG, Yacoub MH, Baines AJ, et al. Expression of human membrane skeleton protein genes for protein 4.1 and betaIISigma2-spectrin assayed by real-time RT-PCR. Cell Mol Biol Lett 2005;10:135–49.

72 Terada N, Ohno N, Saitoh S, Saitoh Y, Komada M, Kubota H, et al. Involvement of a membrane skeletal protein, 4.1G, for Sertoli/germ cell interaction. Reproduction 2010;139:883–92. 10.1530/REP-10-0005.

73 Yang S, Weng H, Chen L, Guo X, Parra M, Conboy J, et al. Lack of protein 4.1G causes altered expression and localization of the cell adhesion molecule nectin-like 4 in testis and can cause male infertility. Mol Cell Biol 2011;31:2276–86. 10.1128/MCB.01105-10.

74 Smith LB, Milne L, Nelson N, Eddie S, Brown P, Atanassova N, et al. KATNAL1 regulation of sertoli cell microtubule dynamics is essential for spermiogenesis and male fertility. PLoS Genet 2012;8:e1002697. 10.1371/journal.pgen.1002697.

75 Cerván-Martín M, Bossini-Castillo L, Guzmán-Jiménez A, Rivera-Egea R, Garrido N, Lujan S, et al. Common genetic variation in KATNAL1 non-coding regions is involved in the susceptibility to severe phenotypes of male infertility. Andrology 2022;10:1339–50. 10.1111/andr.13221.

76 Tang EI, Cheng CY. MARK2 and MARK4 Regulate Sertoli Cell BTB Dynamics Through Microtubule and Actin Cytoskeletons. Endocrinology 2022;163:bqac130. 10.1210/endocr/bqac130.

77 Windley SP, Mayère C, McGovern AE, Harvey NL, Nef S, Schwarz Q, et al. Loss of NEDD4 causes complete XY gonadal sex reversal in mice. Cell Death Dis 2022;13:75. 10.1038/s41419-022-04519-z.

78 Zenker M, Edouard T, Blair JC, Cappa M. Noonan syndrome: improving recognition and diagnosis. Arch Dis Child 2022;107:1073–8. 10.1136/archdischild-2021-322858.

79 Chohan H, Esfandiarei M, Arman D, Van Raamsdonk CD, van Breemen C, Friedman JM, et al. Neurofibromin haploinsufficiency results in altered spermatogenesis in a mouse model of neurofibromatosis type 1. PLoS One 2018;13:e0208835. 10.1371/journal.pone.0208835.

80 Feng Y, Chen D, Wang T, Zhou J, Xu W, Xiong H, et al. Sertoli cell survival and barrier function are regulated by miR-181c/d-Pafah1b1 axis during mammalian spermatogenesis. Cell Mol Life Sci 2022;79:498. 10.1007/s00018-022-04521-w.

81 Dickins RA, Frew IJ, House CM, O’Bryan MK, Holloway AJ, Haviv I, et al. The ubiquitin ligase component Siah1a is required for completion of meiosis I in male mice. Mol Cell Biol 2002;22:2294–303. 10.1128/MCB.22.7.2294-2303.2002.

82 Buratti J, Ji L, Keren B, Lee Y, Booke S, Erdin S, et al. De novo variants in SIAH1, encoding an E3 ubiquitin ligase, are associated with developmental delay, hypotonia and dysmorphic features. J Med Genet 2021;58:205–12. 10.1136/jmedgenet-2019-106335.

83 Whitmill A, Timani KA, Liu Y, He JJ. Tip110: Physical properties, primary structure, and biological functions. Life Sci 2016;149:79–95. 10.1016/j.lfs.2016.02.062.

84 Ayers KL, Eggers S, Rollo BN, Smith KR, Davidson NM, Siddall NA, et al. Variants in SART3 cause a spliceosomopathy characterised by failure of testis development and neuronal defects. Nat Commun 2023;14:3403. 10.1038/s41467-023-39040-0.

